# Structure insights, thermodynamic profiles, dsDNA melting activity, and liquid-liquid phase separation of the SARS-CoV-2 nucleocapsid N-terminal domain binding to DNA

**DOI:** 10.1101/2021.07.21.453232

**Authors:** Icaro Putinhon Caruso, Vitor dos Santos Almeida, Mariana Juliani do Amaral, Guilherme Caldas de Andrade, Gabriela Rocha de Araújo, Talita Stelling de Araújo, Jéssica Moreira de Azevedo, Glauce Moreno Barbosa, Leonardo Bartkevihi, Peter Reis Bezerra, Katia Maria dos Santos Cabral, Isabella Otênio Lourenço, Clara L. F. Malizia-Motta, Aline de Luna Marques, Nathane Cunha Mebus-Antunes, Thais Cristtina Neves-Martins, Jéssica Maróstica de Sá, Karoline Sanches, Marcos Caique Santana-Silva, Ariana Azevedo Vasconcelos, Marcius da Silva Almeida, Gisele Cardoso de Amorim, Cristiane Dinis Anobom, Andrea T. Da Poian, Francisco Gomes-Neto, Anderson S. Pinheiro, Fabio C. L. Almeida

**Affiliations:** Institute of Medical Biochemistry, Federal University of Rio de Janeiro, Rio de Janeiro, Brazil; Multiuser Center for Biomolecular Innovation (CMIB), Department of Physics, São Paulo State University (UNESP), São José do Rio Preto, Brazil; National Center of Nuclear Magnetic Resonance (CNRMN), CENABIO, Federal University of Rio de Janeiro, Rio de Janeiro, Brazil; Faculty of Pharmacy, Federal University of Rio de Janeiro, Rio de Janeiro, Brazil; Protein Advanced Biochemistry (PAB), CENABIO, Federal University of Rio de Janeiro, Rio de Janeiro, Brazil; Department of Biochemistry, Institute of Chemistry, Federal University of Rio de Janeiro, Rio de Janeiro, Brazil; Multidisciplinary Center for Research in Biology (NUMPEX), Campus Duque de Caxias Federal University of Rio de Janeiro, Duque de Caxias, Brazil; Laboratory of Toxinology, Oswaldo Cruz Foundation (FIOCRUZ), Rio de Janeiro, Brazil; Rio BioNMR Network, Rio de Janeiro, Brazil

**Keywords:** SARS-CoV-2, nucleocapsid protein, TRS, DNA/RNA binding protein, melting activity, liquid-liquid phase separation

## Abstract

The SARS-CoV-2 nucleocapsid protein (N) is a multifunctional promiscuous nucleic acid-binding protein, which plays a major role in nucleocapsid assembly and discontinuous RNA transcription, facilitating the template switch of transcriptional regulatory sequences (TRS). Here, we dissect the structural features of the N protein N-terminal domain (N-NTD), either with or without the SR-rich motif (SR), upon binding to single and double-stranded TRS DNA, as well as their activities for dsTRS melting and TRS-induced liquid-liquid phase separation (LLPS). Our study gives insights on specificity for N-NTD/N-NTD-SR interaction with TRS, including an unfavorable energetic contribution to binding along with hydrogen bonds between the triple-thymidine (TTT) motif in the dsTRS and β-sheet II due to the defined position and orientation of the DNA duplex, a well-defined pattern (ΔH > 0 and ΔS > 0 for ssTRS, and ΔH < 0 and ΔS < 0 for dsTRS) for the thermodynamic profile of binding, and a preference for TRS in the formation of liquid condensates when compared to a non-specific sequence. Moreover, our results on DNA binding may serve as a starting point for the design of inhibitors, including aptamers, against N, a possible therapeutic target essential for the virus infectivity.

## INTRODUCTION

The pandemic of severe acute respiratory syndrome coronavirus 2 (SARS-CoV-2), the causative agent of Covid-19, is still a global health emergency. SARS-CoV-2 is an airborne virus that has rapidly spread around the world, infecting more than 80 million people, and causing almost 2 million deaths. The infection in humans is mainly through inhalation (1, 2) and the typical clinical symptoms include fever, cough, sore throat, fatigue, headache, myalgia, and breathlessness. It can progress to severe pneumonia, respiratory failure, and can be fatal (1, 3).

Coronaviruses (CoVs) are enveloped viruses containing a single-stranded positive-sense large genome composed of up to 32 kb. CoV genome replication is a continuous process, whereas transcription requires a complex discontinuous mechanism that culminates in the synthesis of a nested set of subgenomic mRNAs (sgmRNAs) (4). This discontinuous transcription process is controlled by a specific sequence motif called transcriptional regulatory sequences (TRS) (5). The common leader TRS (TRS-L) is present at the 5’-untranslated region of the genome, while other TRSs containing conserved core sequences are located upstream of each ORF, named body TRSs (TRS-B). During RNA synthesis, the negative-sense RNA strand containing TRS-B is transferred to and hybridizes with TRS-L. This allows the RNA-dependent RNA polymerase to switch template to TRS-L, generating a set of short negative-sense RNAs that serve as a template for the transcription of positive-sense sgmRNAs (4). sgmRNAs code for several accessory and structural proteins, including spike (S), membrane (M), envelope (E), and nucleocapsid (N). Recently, subgenomic RNA transcription has been validated for SARS-CoV-2 (6).

N protein is encoded by the most abundant sgmRNA during infection. It is a multifunctional RNA-binding protein that plays critical roles in nucleocapsid assembly as well as in the regulation of RNA replication, transcription, and translation (7). Moreover, it is the only viral structural protein found at the replicase-transcriptase complex (8). N is a protein dimer composed of two globular domains, an N-terminal domain (NTD) and a C-terminal domain (CTD), which are flanked by flexible regions, including the N-terminal arm, the central Ser/Arg rich (SR) intrinsically disordered linker named LKR, and the C-terminal tail (7). The CTD mediates N protein dimerization (9, 10). In addition, the NTD, the CTD, and the intrinsically disordered regions all engage in RNA binding (11). N protein has been shown to participate in CoVs’ subgenomic RNA transcription. Deletion of N protein from the replicon significantly reduces the levels of sgmRNA (12). Furthermore, N-NTD specifically interacts with TRS and efficiently melts double-stranded TRS RNA. This RNA chaperone activity is suggested to facilitate template switch and thus regulate discontinuous transcription (10, 13–15). However, the structural basis for TRS binding specificity and the molecular mechanism by which N-NTD melts dsRNA are still poorly understood.

Multivalent RNA-binding proteins, especially those containing intrinsically disordered regions and low complexity domains, have been shown to undergo liquid-liquid phase separation (LLPS) (16). LLPS is the physical principle underlying the formation of membraneless organelles, dynamic micron-sized compartments that provide spatiotemporal control of cellular reactions and organization of biomolecules (17, 18). Interestingly, LLPS has been regarded as an universal nucleation step for the assembly of several viruses, including Measles (19). Previous reports have demonstrated that SARS-CoV-2 N forms liquid condensates upon RNA interaction (20–23). RNA-driven LLPS of N is mainly mediated by electrostatic interactions (21) and is regulated by RNA sequence and structure (22). Remarkably, N liquid droplets recruit and concentrate components of the replication machinery (21). Despite these recent findings, the role played by TRS binding on N condensation remains unexplored.

Here, we report on the ability of N-NTD/N-NTD-SR to melt dsTRS DNA and to undergo LLPS upon TRS binding. Critically, N-NTD-SR droplet formation was dependent on the intrinsically disordered SR-rich region. Based on experimental NMR data, we calculated structural models for the interaction of N-NTD with single and double-stranded TRS DNAs. DNA oligonucleotides bind at the positively charged cleft between the palm and the finger, similarly to RNA (15). Electrostatic interactions play a major role in complex stabilization, specifically by protein-DNA hydrogen bonds and salt bridges. dsTRS binding is enthalpically favorable and entropically unfavorable, while the opposite is observed for ssTRSs. We propose that TRS binding leads to a well-defined positioning of the nucleic acid duplex with a geometry favorable for melting activity. This positioning finely tunes favorable and unfavorable contributions to the binding free energy that may balance the biological activity and explain specificity.

## MATERIAL AND METHODS

### Protein expression and purification

The gene sequences encoding the SARS-CoV-2 (NC_045512.2) nucleocapsid protein N-terminal domain (N-NTD; residues 44–180) and the N-NTD containing the C-terminal SR-rich motif (N-NTD-SR; residues 44–212) were codon-optimized, synthesized, and subcloned into pET28a by GenScript (Piscataway, USA). Expression and purification of these constructs were previously described by the international Covid19-NMR consortium (https://covid19-nmr.de/) (24). Briefly, pET28a-N-NTD and pET28a-N-NTD-SR were transformed into *Escherichia coli* BL21 (DE3). Cells were grown at 37 °C until optical density ∼0.7 at 600 nm. Protein expression was induced with 0.2 mM IPTG for 16 h at 16 °C. For the production of ^15^N and ^15^N/^13^C-labeled protein, expression was induced in M9 minimal medium containing either ^15^NH_4_Cl (1.0 g/L) or ^15^NH_4_Cl (1.0 g/L) and ^13^C-glucose (3.0 g/L). All media contained 50 µg/mL of kanamycin. Cells were harvested by centrifugation (10,000 rpm, 4 °C, 15 min), resuspended in 30 mL of lysis buffer containing 50 mM Tris-HCl (pH 8.0), 500 mM NaCl, 20 mM imidazole, 10% glycerol, 0.01 mg/mL DNAase, 5 mM MgCl, and 0.1 mM PMSF, and lysed by sonication. The supernatants were clarified by centrifugation (10,000 rpm, 4 °C, 15 min) and subsequently applied to a HisTrap FF column (Cytiva) and purified by nickel-affinity chromatography. The washing buffer contained 50 mM Tris-HCl (pH 8.0), 500 mM NaCl, 20 mM imidazole, and 10% glycerol. The bound proteins were eluted with a linear 20–500 mM imidazole gradient. Fractions containing the proteins of interest were pooled and dialyzed overnight at 4 °C against buffer containing 50 mM Tris-HCl (pH 8.0), 500 mM NaCl, and 1.0 mM DTT. Simultaneously, the constructs were cleaved with TEV protease (1:30 TEV:protein molar ratio) to remove the His_6_ tag. After dialysis, samples were reapplied to a HisTrap FF column, using the same purification buffers. Fractions containing the proteins of interest were pooled, dialyzed against buffer containing 20 mM sodium phosphate (pH 6.5) and 50 mM NaCl, and concentrated. Protein concentration was determined spectrophotometrically using the molar extinction coefficient of 26,930 M^−1^·cm^−1^ at 280 nm.

### DNA fragments

In this study, we used a 11-nucleotide DNA sequence corresponding to a 7-nucleotide conserved sequence of the positive-sense ssTRS followed by a CGCG generic segment (ssTRS(+), 5’–TCTAAACCGCG–3’). This CGCG sequence increased the melting temperature of the corresponding TRS duplex and thus enabled us to work at room temperature. The negative-sense ssTRS contained the complementary strand (ssTRS(–), 5’–CGCGGTTTAGA–3’). As a control, we used the single and double-stranded non-specific (NS) oligonucleotides containing the 7-nucleotide DNA sequences used by Dinesh and cols. (2020) (15) followed by the CGCG generic segment (ssNS(+), 5’–CACTGACCGCG–3’; and ssNS(–), 5’–CGCGGTCAGTG–3’). All DNA oligonucleotides were purchased from GenScript (Piscataway, USA). Equimolar amounts of positive and negative sense ssDNAs were dissolved in buffer and annealed by heating to 50 °C for 10 min and slowly cooling to room temperature. The concentrations of ssTRS(+), ssTRS(–), ssNS(+), and ssNS(–) were determined spectrophotometrically using the molar extinction coefficients of 103000, 107500, 99100, and 103000 M^−1^·cm^−1^ at 260 nm, respectively.

### Fluorescence spectroscopy

Protein intrinsic fluorescence quenching measurements were performed using a PC1 steady-state spectrofluorimeter (ISS, Champaign, IL, USA) equipped with a quartz cell of 1.0 cm optical path length and a Neslab RTE-221 thermostat bath. Excitation and emission bandwidths were set to 1.0 and 2.0 mm, respectively. To solely excite tryptophan, the excitation wavelength (*λ_ex_*) was set to 295 nm. The emission spectrum was collected in the range of 305–500 nm with an increment of 1.0 nm. Each point in the emission spectrum is the average of 10 accumulations. Increasing concentrations of DNA oligonucleotides were added to a 1.0 μM protein solution in 20 mM Bis-Tris buffer (pH 6.5). The DNA concentration varied from 0 to 3.8 μM, and experiments were performed in either duplicate or triplicate. Titration measurements were performed at 15, 25, and 35 °C for ssDNAs, and 15, 20, and 25 °C for dsDNAs. The effect of salt (NaCl) and inorganic phosphate (Pi) on the N-NTD:ssDNA complex formation was investigated at 25 °C. The binding isotherms were fitted using the following equation after data treatment (25):

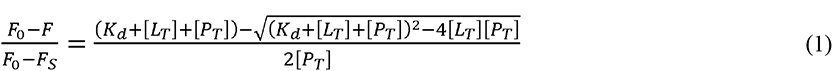

where *F* is the measured fluorescence intensity in the presence of DNA, *F_0_* is the fluorescence in the absence of DNA, *F_S_* is the fluorescence emission of the protein saturated with DNA, *K_d_* is the dissociation constant, [*L_T_*] is the total ligand concentration (DNA), and [*P_T_*] is the total protein concentration (N-NTD or N-NTD-SR). The *K_d_* values were calculated from the fitting process by nonlinear least-squares optimization using Levenberg–Marquardt interactions in the software OriginPro 2021. The global fitting was performed for the protein:DNA binding isotherms at three temperatures using as a constraint the linearity of the van’t Hoff plot, *ln*(*K_d_*) versus *T^−1^*, where *T* is the temperature in Kelvin. The linear constraint is based on the non-significant variation of the enthalpy change (Δ*H*) in the temperature range studied (15–35 °C for ssDNAs, and 15–25 °C for dsDNAs). Next, Δ*H* values were determined from the van’t Hoff equation:

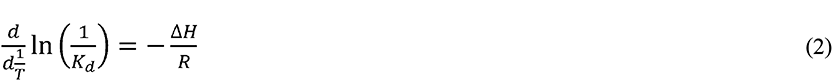

where R is the universal gas constant. The values of the Gibbs free energy changes (Δ*G*) and entropy change (Δ*S*) were determined from Δ*G* = *RTln*(*K_d_*) and TΔ*S* = ΔH - ΔG.

Prior to fitting, the fluorescence quenching data were corrected for the background buffer fluorescence, the inner filter effects (26), and for removing an upward linear contribution from the fluorescence titration curves at the largest DNA concentrations (Figure S1). This near-linear contribution arises due to the difference in the accessibility of the fluorophores (W52, W108, and W132) by the quencher (DNAs). Two of the three fluorophores (W52 and W108) are located at the putative nucleic acid-binding site in N-NTD, while the third (W132) is oppositely oriented to this binding site and thus presents a non-specific linear contribution for the fluorescence titration curves. To remove the non-specific contribution, *F_0_*/*F versus* [*L_T_*] curves for each protein:DNA titration (duplicate or triplicate) was fitted using Equation (1) plus a linear term (*constant*·[*L_T_*]). After that, *F_0_*/*F* linear contributions were calculated and removed manually (Figure S1).

Fluorescence resonance energy transfer (FRET) experiments were performed on a Cary Eclipse spectrofluorometer (Agilent Technologies, Santa Clara, CA, USA) equipped with a quartz cell of 1.0 cm optical path length and Peltier temperature controller at 20 °C. A bandwidth of 5 nm was set for both excitation and emission. FRET measurements were carried out with DNA and RNA oligonucleotides labeled at the 5’-end of the positive-sense TRS strand with the donor probe Q570 (Q570-ssTRS(+)) and at the 3’-end of the negative-sense TRS strand with the acceptor probe Q670 (Q670-ssTRS(–)). The donor Q570 was excited at 535 nm and the emission spectrum was collected in the 550–800 nm range, observing the FRET efficiency (*E*) of the acceptor Q670 at maximum emission at 670 nm. The oligonucleotide concentration was set to 50 nM, while the final concentrations of protein was in the range of 9–17 μM. Fluorescence measurements were recorded in 20 mM Bis-Tris buffer (pH 6.5) in the absence or presence of 100 mM NaCl, and in 20 mM sodium phosphate buffer (pH 6.5) containing 50 and 100 mM NaCl. The FRET efficiency was calculated from E = 1 – *F_DA_*/*F_D_*, where *F_D_* and *F_DA_* are the fluorescence intensities of the donor Q570 in the absence and presence of the acceptor Q670, respectively.

### NMR spectroscopy

Chemical shift perturbations (CSP) were monitored by a series of 2D [^1^H,^15^N] HSQC spectra, recorded on a Bruker 600 MHz spectrometer at 20 °C, after the addition of increasing concentrations of ssDNAs (TRS and NS). N-NTD and N-NTD-SR were dissolved in 20 mM sodium phosphate buffer (pH 6.5), 50 mM NaCl, 500 µM PMSF, 3 mM sodium azide, 3 mM EDTA, and 5.5% (v/v) of D_2_O, at a final concentration of 70 µM. The ssDNAs were titrated into N-NTD and N-NTD-SR for the following ssDNA:protein molar ratios: 0.14, 0.34, 0.77, 1.00, 1.57, 2.49, and 2.99. For dsDNAs (280 μM), CSPs were calculated from a protein sample containing 4× molar excess of dsDNA. Spectra were processed with NMRPipe (27), and analyzed with CCPNMR Analysis (28). The chemical shift of water was used as an internal reference for ^1^H, while the ^15^N chemical shift was referenced indirectly to water (Wishart et al. 1995). Three [^1^H,^15^N] HSQC spectra were collected for the free protein and used to determine the experimental error. The chemical shift perturbations (CSP, Δδ) of all assigned amides were calculated by:

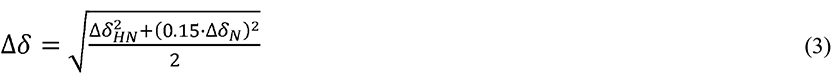

where Δδ*_HN_* and Δδ*_N_* are the chemical shift differences of ^1^H and ^15^N, respectively, recorded in the absence and presence of DNA. The standard deviation of the average weighted Δδ values for all assigned amides was used as a cutoff to identify the protein residues involved in DNA binding. Residues showing fast exchange in the presence of DNA were used for the calculation of *K_d_* from Equation (1).

### Computation simulations

The HADDOCK (version 2.4) server (29) was used to construct a structural model of the N-NTD complex with TRS and NS DNAs. The protein structural coordinates used as input were obtained from the Protein Data Bank (PDB) under access code 6YI3 (15). The ^1^H, ^15^N chemical shifts of residues forming the DNA-binding interface (identified by the CSP analysis) were used as ambiguous restrains for the docking. Residues with chemical shift perturbation Δδ > Δδ*_av_* +2·*SD* were classified as active and those with Δδ > Δδ*_av_* + *SD* were defined as passives. In total, 2000 complex structures of rigid-body docking were calculated by using the standard HADDOCK protocol with an optimized potential for liquid simulation (OPLSX) parameters (30). The final 200 lowest-energy structures were selected for subsequent explicit solvent (water) and semi-flexible simulated annealing refinement, to optimize side chain constants. The final structures were clustered using the fraction of common contacts (FCC) with a cutoff of 0.6 (31).

Molecular dynamics (MD) calculations for the N-NTD:DNA complexes were performed using GROMACS (version 5.1.4) (32). The molecular systems were modeled with the corrected AMBER14-OL15 package, including the ff14sb protein (33) and ff99bsc0OL15 DNA (34) force fields, as well as the TIP3P water model (35). The structural models of the N-NTD:DNA complexes (from molecular docking) were placed in the center of a cubic box solvated by a solution of 50 mM NaCl in water. The protonation state of ionizable residues at pH 7.0 was set according to the PROPKA server (36). Periodic boundary conditions were used, and all simulations were performed in NPT ensemble, keeping the system at 25 °C and 1.0 bar using Nose-Hoover thermostat (τ_T_ = 2 ps) and Parrinello-Rahman barostat (τ_P_ = 2 ps and compressibility = 4.5×10^−5^·bar^−1^). A cutoff of 12 Å for both Lennard-Jones and Coulomb potentials was used. The long-range electrostatic interactions were calculated using the particle mesh Ewald (PME) algorithm. In every MD simulation, a time step of 2.0 fs was used and all covalent bonds involving hydrogen atoms were constrained to their equilibrium distance. A conjugate gradient minimization algorithm was used to relax the superposition of atoms generated in the box construction process. Energy minimizations were carried out with the steepest descent integrator and conjugate gradient algorithm, using 1,000 kJ·mol^−1^·nm^−1^ as maximum force criterion. One hundred thousand steps of molecular dynamics were performed for each NVT and NPT equilibration, applying force constants of 1,000 kJ·mol^−1^·nm^−2^ to all heavy atoms of N-NTD:DNA complexes. At the end of preparation, 2 μs MD simulations of each molecular system (N-NTD bound to ssTRSs, dsTRS, ssNS, and dsNS) were carried out for data acquisition. Following dynamics, the trajectories of each molecular system were firstly concatenated and analyzed according to the root mean square deviation (RMSD) for the backbone atoms of protein and DNA, number of contacts for distances lower than 0.6 nm between pairs of atoms of N-NTD and DNA, and number of protein-DNA hydrogen bonds with cutoff distance (heavy atoms) of 3.5 Å and maximum angle of 30°. The percentages of protein-DNA hydrogen bond persistence were obtained from *plot_hbmap_generic.pl* script (37). The number of protein-DNA hydrogen bonds with persistence greater than 10% was counted with respect to amino acid and nucleotide residues for each molecular system. The contributions of N-NTD residues to the DNA-binding energy were calculated from the MD trajectories using the Molecular Mechanics Poisson-Boltzmann Surface Area (MM-PBSA) method implemented in the *g_mmpbsa* program, along with the *MmPbSaDecomp.py* script (38, 39). The tool *g_cluster* of GROMACS package (40) was used to perform a cluster analysis in the 2.0 μs MD trajectories of the N-NTD:DNA complexes. The *gromos* algorithm (41) was used to generate the clusters with cutoff of 3 and 4 Å for ssDNA and dsDNA, respectively. The RMSD was calculated between all the structures and superimposed considering only non-hydrogen atoms. The principal component analysis (PCA) was applied to investigated conformational space of the protein in the interaction with DNA. PCA scatter plots were generated, and conformational motions were filtered from the eigenvectors of the first and second principal components (PC1 and PC2, respectively). The conformational space was quantified by fitting an elliptical shell with 95% (confidence) of the density for each scatter plot and making its extent proportional to the area (*S_el_*) of this shell. The structural representations of the constructed models were displayed using PyMOL (42).

### Microscopy and turbidity for LLPS study

Cleaning of microscopy glasses followed by coating of Knittel coverslips with 0.5% (w/v) bovine serum albumin (BSA, Sigma-Aldrich A2153) was carried out to provide a hydrophilic surface. Specifically, coverslips were incubated for 1 h with the BSA solution and rinsed twice with Milli-Q water. Subsequently, 5 × 22 mm imaging chambers were mounted onto glass slides using double-sided tapes to stick coverslips as reported in (43, 44). The protein samples of N-NTD and N-NTD-SR at 20 µM were prepared in low-binding microtubes (Eppendorf® LoBind) containing 20 mM Tris-HCl buffer (pH 7.5) with 30 mM NaCl, specified PEG-4000 concentrations (Thermo Scientific, USA), and/or reactants such as 10% (w/v) 1,6-hexanediol (Merck, Germany), 300 mM NaCl. The following nucleic acids were used: a pool of RNA consisting of 5000-8000 Da from Torula yeast (Sigma-Aldrich, R6625), ssTRS(+), ssTRS(–), ssNS(+), and ssNS(–) with concentrations ranging from 2.5 to 40 µM according to the protein:nucleic acid molar ratios (8:1, 4:1, 2:1, 1:1, and 1:2). The dsTRS and dsNS were produced as mentioned above. Immediately following nucleic acid addition to N-NTD solution, a 15 μL aliquot was pipetted through one opening of the rectangular chamber and settle to stand for 30 minutes with coverslip surface facing down to let protein condensates stand by gravity followed by imaging and concomitant turbidity measurement. All samples were imaged at time 30 minutes at room temperature. The working concentration of the fluorescent dye DAPI (BioRad, #135-1303, stock in Milli-Q water) was 0.25 μg/mL and the emission recorded using a LED light fluorescent cube (excitation centered at 357/44 and emission at 447/60 nm). Samples described above were imaged in an inverted phase contrast microscope (EVOS® FL Cell Imaging System, Thermo Fisher Scientific, USA) with a 40× apo objective. To quantify the number of condensates, five areas of 100 µm^2^ were analyzed for each sample followed by Fourier bandpass filtering. Next, images had their threshold adjusted for liquid droplets correct recognition, followed by mask creation and the “fill holes” (fills droplets that were poorly delimited) and “watershed” operations (separate droplets undergoing fusion). The condensates number (above 0.5 µm condensates were counted) and size (area in µm^2^) were determined using the “analyze particles” plugin. All quantification and image processing steps were performed in Fiji. The DAPI fluorescence images were obtained with 1% excitation intensity, apart from the crystal sample formed by 1:1 N-NTD-SR:dsNS which was excited with 20% laser power. Representative micrographs from three independent experiments had their brightness/contrast corrected by histogram stretching. Data were analyzed with GraphPad Prism (version 8.1.1; GraphPad Software).

Turbidity (detected as absorbance at 350 nm) was performed concomitant to imaging time (30 min). Samples of 20 μΜ N-NTD or N-NTD-SR consisting of varied protein:nucleic acid molar ratios (8:1, 4:1, 2:1, 1:1, 1:2) were prepared in low-binding microtubes (Eppendorf® LoBind) in the presence of 10% PEG-4000 (w/v) in 20 mM Tris-HCl buffer (pH 7.5) with 30 mM NaCl. Three technical replicates were measured using 5 µL of samples in a μLite sample port (0.5 mm optical path) of a BioDrop Duo spectrophotometer (Biochrom, UK). Protein:nucleic acid sample readings were subtracted from samples containing all components apart from protein. Data were analyzed with GraphPad Prism (version 8.1.1; GraphPad Software).

## RESULTS

### N-NTD and N-NTD-SR melt double-stranded DNA

We first measured the dsDNA melting activity of N-NTD, as well as N-NTD containing the SR-rich motif (N-NTD-SR), using fluorescence resonance energy transfer (FRET) from the 5’ Q570-labeled positive-sense TRS (Q570-ssTRS(+)) to the 3’ Q670-labeled negative-sense TRS (Q670-ssTRS(–)) (Figure 1A). The DNA was used as a N-NTD/N-NTD-SR ligand due to its higher stability over RNA, while maintaining a relative structural similarity. As it will be further described, the use of DNA is supported by the chemical shift profiles observed for dsDNA (either TRS or NS), which are largely similar to those obtained with dsRNA (9). The study with DNA is complementary to the available data on the N protein interaction with RNA (7, 15, 45). In the absence of N-NTD, the FRET pair Q570/Q670 is in proximity, displaying FRET efficiency for dsTRS. Addition of N-NTD or N-NTD-SR results in an increase in fluorescence emission at 570 nm with a concomitant decrease at 670 nm, leading to a decay in FRET efficiency, indicating DNA duplex melting (Figure 1). N-NTD and N-NTD-SR shows similar dsTRS melting activity. We also measured N-NTD-SR melting activity toward dsTRS RNA, which presented similar activity (Figure S2).

**Figure 1.**
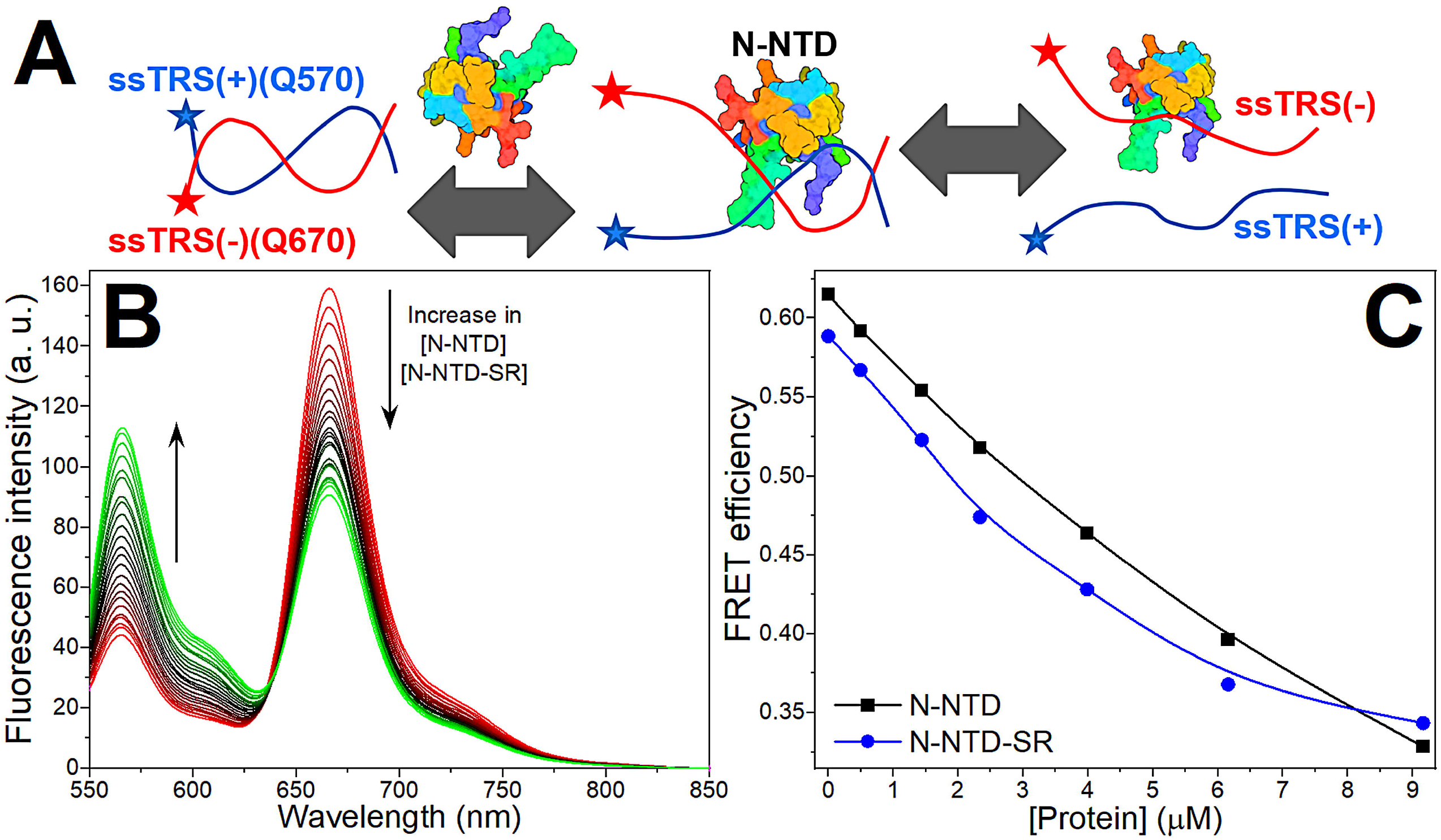
dsDNA melting activity of N-NTD and N-NTD-SR. (A) Cartoon model of the dsTRS melting activity of N-NTD(-SR). The ssTRSs are presented as curved lines and the fluorescent probes (Q570 and Q670) as stars. ssTRS(+) and ssTRS(–) are colored in blue and red, respectively. The protein is denoted as a colored surface. (B) Fluorescence spectra of the Q570-ssTRS(+)/Q670-ssTRS(–) DNA duplex (50 nM) at absence and presence of N-NTD or N-NTD-SR (*λ_ex_* = 535 nm and temperature at 20 °C). The arrows denote the changes in the spectra as protein is added. (C) FRET efficiency as a function of the total concentration of N-NTD (black squares) and N-NTD-SR in (blue circles) 20 mM sodium phosphate buffer (pH 6.5) containing 50 mM NaCl.

### N-NTD and N-NTD-SR bind tighter to ssTRS(–) than ssTRS(+)

We investigated the thermodynamics of N-NTD/N-NTD-SR binding to the specific (ssTRS(+), ssTRS(–), and dsTRS) and non-specific DNA sequences (ssNS(+), ssNS(–), and dsNS), using intrinsic tryptophan fluorescence. Using 20 mM Bis-Tris buffer, titration of DNA oligonucleotides into a protein solution resulted in fluorescence quenching until saturation, indicating protein:DNA association. After data treatment, the binding isotherms exhibited a hyperbolic shape (Figure 2A and S3). Formation of N-NTD:DNA complexes showed salt dependence. We observed a fluorescence recovery upon the increase of inorganic phosphate (Pi) and sodium chloride (NaCl) concentrations (Figure 2B and 2C), suggesting that electrostatic interactions are essential for complex formation. Despite the high salt and phosphate dependence, N-NTD-SR remains active, independently of the buffer (20 mM Bis-Tris or 20 mM NaPi) and salt concentration (0, 50, or 100 mM NaCl) (Figure S2).

**Figure 2.**
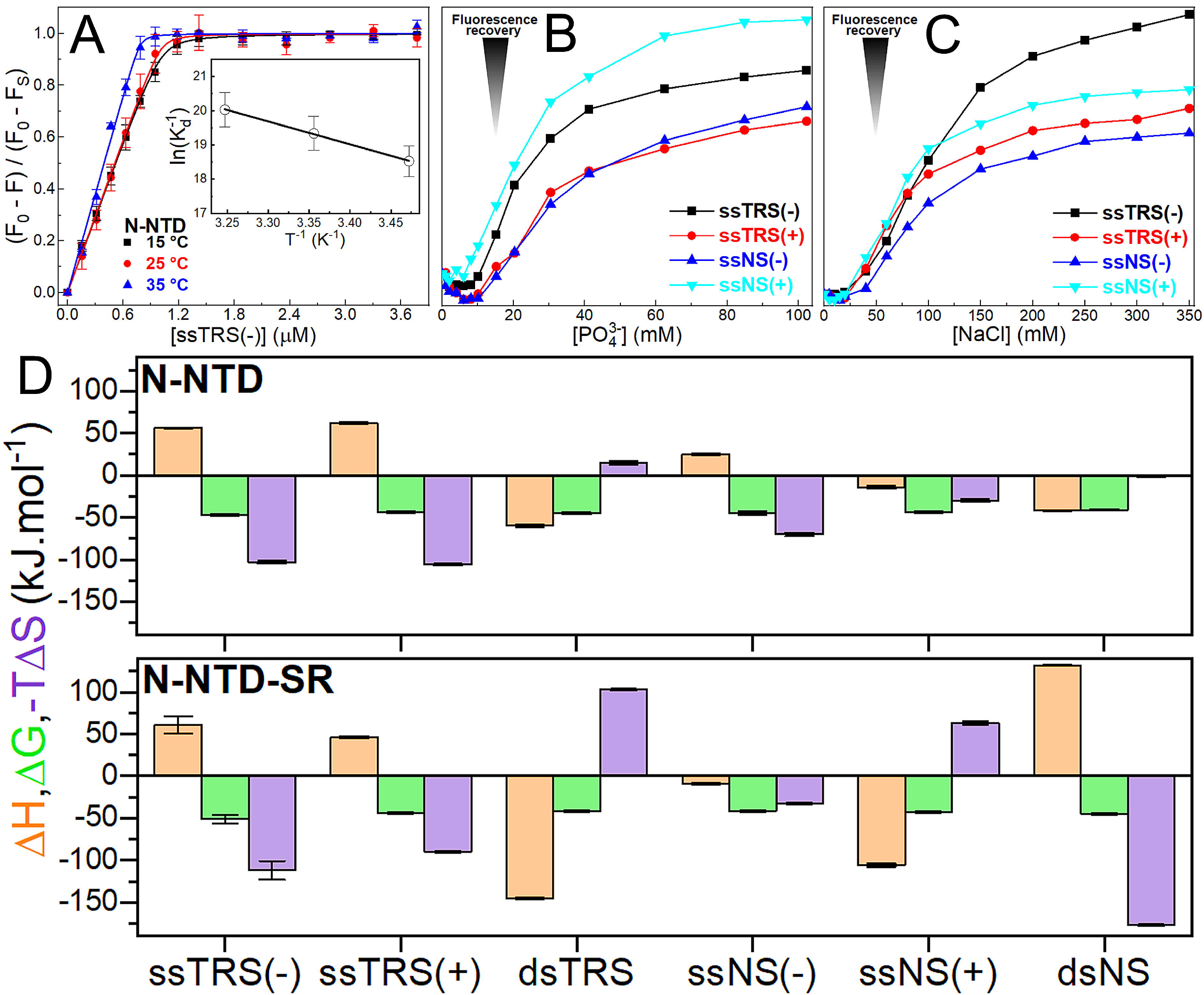
Thermodynamic analysis of the binding of TRS and NS DNAs to N-NTD and N-NTD-SR. (A) Protein intrinsic fluorescence quenching changes of N-NTD as a function of the ssTRS(–) concentration in 20 mM Bis-Tris buffer (pH 6.5) at 15, 25, and 35°C. Each point on the binding isotherm represents the average and standard error calculated from triplicate measurements. The continuous lines denote the theoretical curves globally adjusted to the experimental data. The inset shows the van’t Hoff plot determined the enthalpy change value for the N-NTD/ssTRS(–) complex. (B, C) Fluorescence recovery as a function of inorganic phosphate (Pi) and sodium chloride (NaCl) concentration for the formation of the N-NTD:ssDNA complexes. (D) Thermodynamic parameters for the interaction of TRS and NS DNAs with the N-NTD and N-NTD-SR at 25 °C. For each DNA, the orange bar on the left denotes the enthalpy change (Δ*H*); the green bar in the middle, the Gibbs free energy change (Δ*G*); and the purple on the right, the entropic term (*T*Δ*S*). The error bars represent the standard deviation calculated from duplicate or triplicate measurements.

The protein:DNA binding affinities are at the nanomolar range (Table S7). The affinity for ssTRS(–) is higher than that for ssTRS(+), being 4–5-fold for N-NTD and 15–16-fold for N-NTD-SR. This tendency was also observed for the NS DNA, albeit with a smaller magnitude. This observation is remarkable since ssTRS(–) is the strand transferred to TRS-L during template switch (4). The thermodynamic parameters are similar for both N-NTD and N-NTD-SR binding to TRS. For ssTRSs, the interaction is enthalpically unfavorable (ΔH > 0) and entropically driven (ΔS > 0), while for dsTRS, it is enthalpically driven (ΔH < 0) and entropically unfavorable (ΔS < 0). The analysis of these enthalpic and entropic contributions reveals that hydrophobic contacts are important for N-NTD/N-NTD-SR binding to ssTRSs, while hydrogen bonds (and salt bridges) and van der Waals interactions play a major role in stabilizing the interaction with dsTRS (46). Based on this observation, we hypothesize that the destabilization of dsTRS Watson-Crick hydrogen bonds induced by N-NTD or N-NTD-SR binding would be entropically driven. It would also be enthalpically unfavorable, which might contribute to further dissociation of ssTRSs.

In general, we observed a higher affinity for TRS than for NS in the interaction with N-NTD and N-NTD-SR, except for ssTRS(+) (Table S7). For N-NTD-SR, the affinity is even higher, especially for ssTRS(–), suggesting that the arginine residues in the SR-rich motif may contribute to binding with non-specific electrostatic interactions. Unlike the thermodynamic pattern observed for N-NTD/N-NTD-SR binding to the specific sequences (TRSs), we did not observe a regular pattern for the enthalpic and entropic contributions to the interaction with NS DNAs (Figure 2D and Table S1). Although a thermodynamic profile similar to that of TRS was observed for N-NTD binding to NS DNAs, a different energetic profile was observed for N-NTD-SR.

### Mapping the N-NTD and N-NTD-SR interaction with DNA

To map the residues involved in N-NTD and N-NTD-SR interaction with DNA, single and double-stranded DNA oligonucleotides were titrated into ^15^N-labeled protein samples and amide chemical shifts were measured from 2D [^1^H,^15^N] HSQC spectra (Figure S4). Residues displaying statistically significant chemical shift perturbation (CSP) values (higher than the average plus one standard deviation (SD)) are located in well-defined regions (Figure 3 and S5). Using the experimental CSP data, we modeled the interaction of N-NTD with the DNAs (Figures 3G-3I). The structural models were fundamental to discriminate the interacting amino acid residues from the ones that seem to undergo allosteric effects upon DNA binding. The binding site includes: (*i*) the flexible N-terminal region (N47, N48, T49), (*ii*) the palm formed by β-sheet I (β2/β3/β4) (Y86, A89, R107, W108, Y109, Y111, I131, W132) and the N-terminal residues (A50, S51, F53, T54, A55), (*iii*) the finger (β2/β3 loop) (A90, T91, R92, R93, I94, R95, G96, K100, K102, D103, L104, S105, R107), (*iv*) the β-sheet II (β1/β5) (L56, T57, Q58, H59, G170, Y172, A173, E1 74), (*v*) the thumb (α2/β5 loop) (R149, N150, A152, A156, I157), and (*vi*) the C-terminal region (G175, S176, R177). We also observed significant CSPs for residues L64, K65, F66, G164, T165, T166, and L167, which are located in a remote region from the principal binding site (Figure 3G and 3H), suggesting that these residues form either a secondary binding site, interacting with DNA directly, or an allosteric site, undergoing indirect conformational changes due to DNA binding. In addition, residues at the SR motif (N192, S193, N196, and S197) displayed significant CSPs, suggesting that this region engages in DNA binding. The observed CSPs are strikingly similar (even for the remote region) to those observed by Dinesh and cols. (2020) for a 7-nucleotide and a 10-nucleotide ssTRS(+) RNA (5’–CUAAACG–3’, 5’–UCUCUAAACG–3’), and a 7-nucleotide non-specific RNA duplex (5’–CACUGAC–3’ and 5’–GUCAGUG–3’) (47), suggesting that DNA and RNA bind at the same interface on N-NTD.

**Figure 3.**
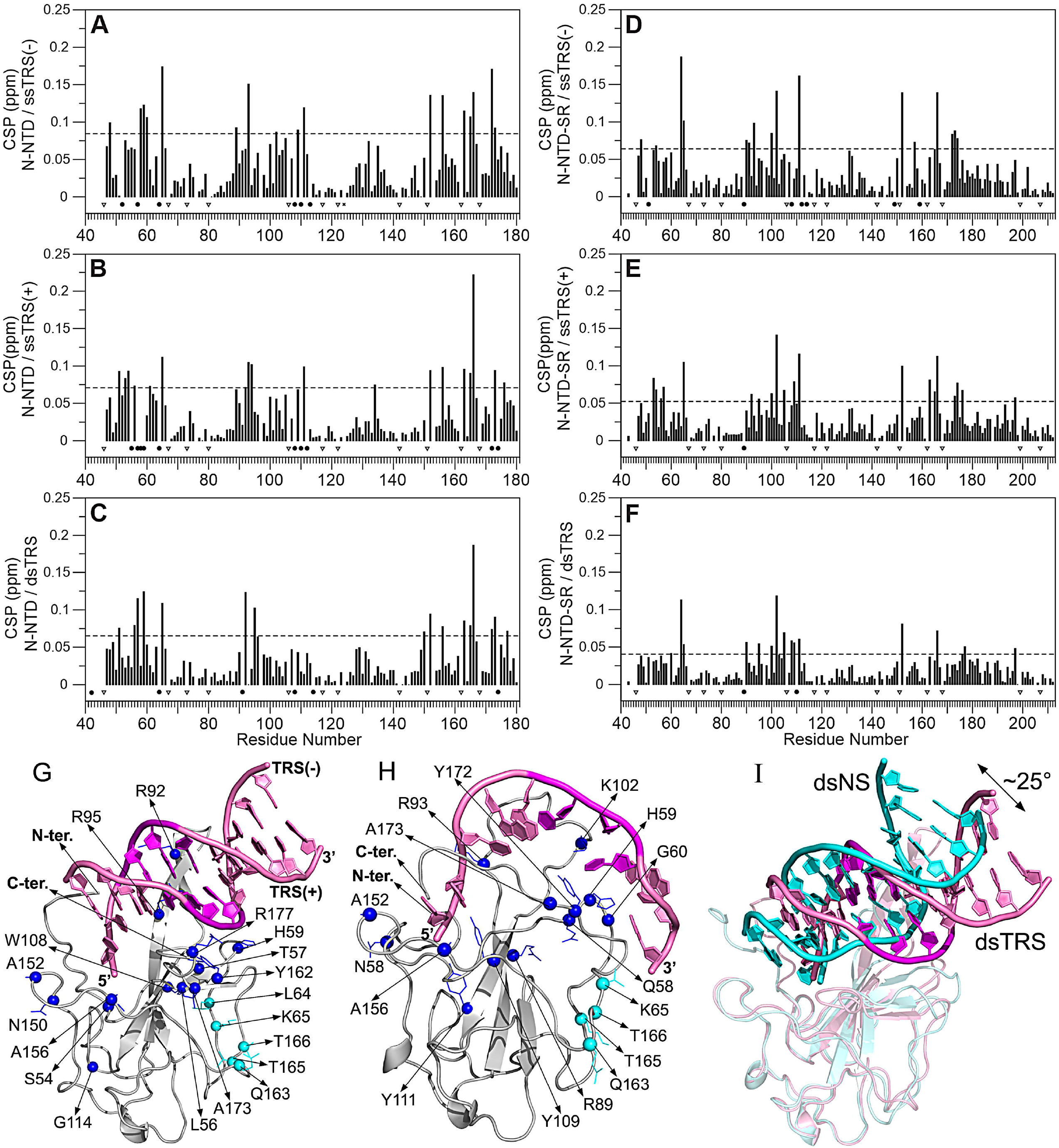
Mapping the residues involved in the protein/TRS interaction. Chemical shift perturbation (CSP) for the interaction of the N-NTD with (A) ssTRS(–), (B) ssTRS(+), and (C) dsTRS, and of the N-NTD-SR with (D) ssTRS(–), (E) ssTRS(+), and (F) dsTRS. The dotted line denotes the average CSP value (Δ*δ_ave_*) plus one standard deviation (SD), which is the cutoff used to identify the most significant residues involved in the binding to DNA. The proline residues (46, 67, 73, 80, 106, 117, 122, 142, 151, 162, 168, 199, and 207) are indicated by triangle. Resonances signals broadened beyond detection upon TRS titration are represented by the filled black circles. Representative structural model of the N-NTD:DNA complexes for (G) dsTRS and (H) ssTRS(–) obtained from the molecular docking and molecular dynamic simulations. The spheres denote the residues with CSP values higher than Δ*δ_ave_* + SD, participating directly in the binding interface (blue) and in the remote region (cyan). (I) Comparison of the position and orientation for dsTRS (light pink) and dsNS (cyan) with a difference of ∼25° between them. The proteins are shown as a cartoon model, DNAs are represented as a cartoon-ring model, and the TTT motif in TRS is colored in magenta.

To find a consensus amino acid sequence for the interaction of N-NTD and N-NTD-SR with ssDNAs and dsDNAs, we aligned the residues with CSPs larger than the average plus one SD for all titrations (Figure S6). To identify which residues are common to all interactions and which are unique, we grouped the different titration results and plotted them as an intersecting set of residues represented by circles containing the CSP information for each titration (Figure 4). When we analyzed the set of intersections for N-NTD and N-NTD-SR with single and duplex TRS, as well as with the non-specific (NS) sequences (Figure 4A, 4B, 4C, and 4D), the following residues stood out: (*i*) A152 in the thumb is present in all 4 intersections; (*ii*) Y111 in the interface between the palm and the β-sheet II, and T166 in the remote region from the main binding site are present in 3 intersections; (*iii*) N47 in the N-terminus, K65 in the remote region, R95 and K102 in the finger, A156 in the thumb, and Y172 and A173 in the β-sheet II are present in 2 intersections. Note that N47 seems to be unique for NS, while K65 for TRS. The regions mapped by the residues at the intersections involve the finger, the two elements of the palm, and the flexible thumb of N-NTD (Figure 4E). These residues may be key for N-NTD binding to several nucleic acids, independent of the sequence specificity. Furthermore, they comprise not only positively charged residues, responsible for electrostatic interactions, but also hydrophobic residues (Y111, A156, T166, Y172, and A173).

**Figure 4.**
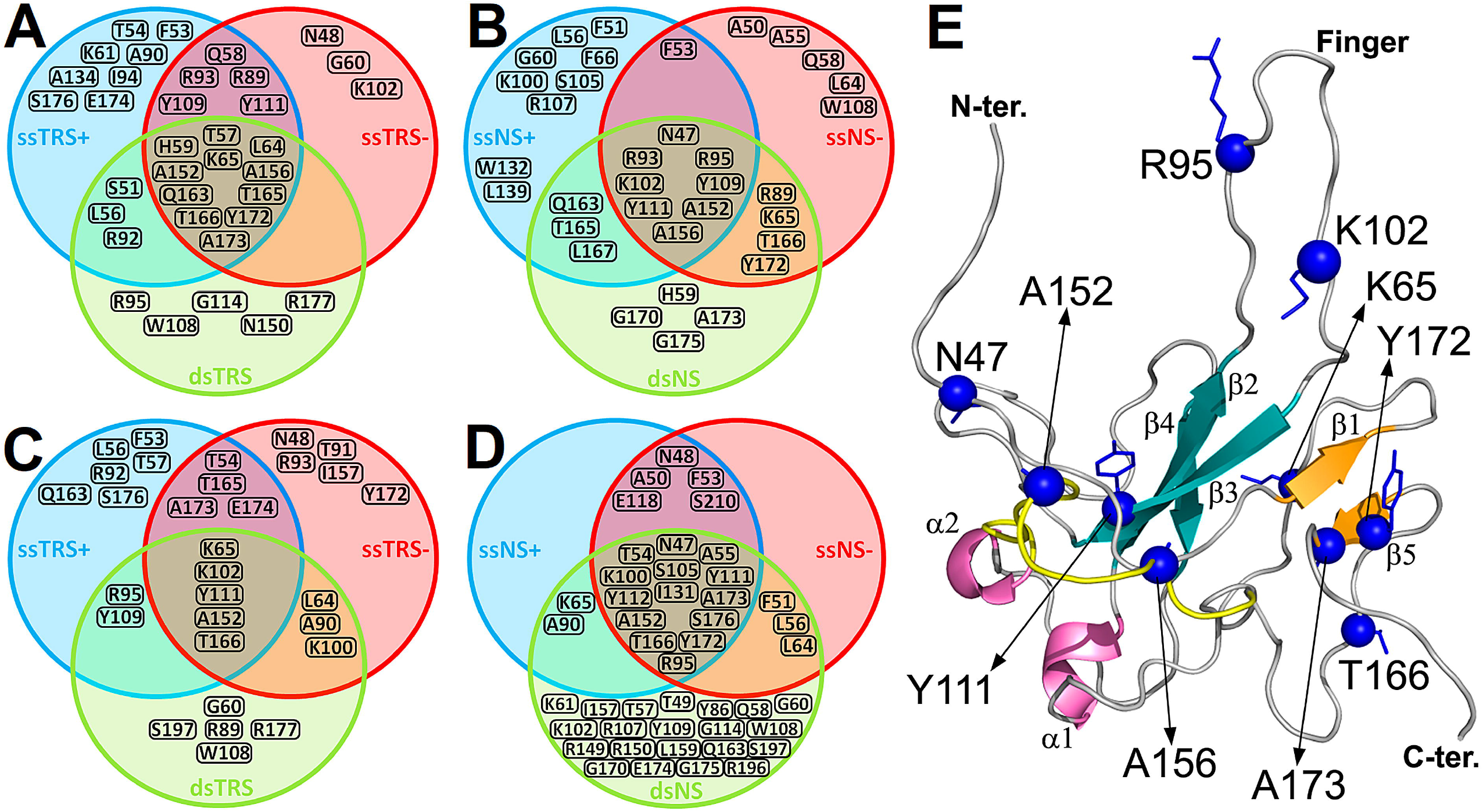
Consensus of residues involved in the protein:DNA binding. Intersecting set of residues represented by circles containing the significant CSP information (higher than Δ*δ_ave_* + SD) for the interaction of the TRS and NS with (A and B) N-NTD and (C and D) N-NTD-SR. The blue, red, and green sphere denotes the CSP set for ssDNA(+), ssDNA(–), and ds DNA, respectively. (E) The DNA-binding residues most recurrent in the intersections for the protein:DNA binding are indicated as blue spheres on the N-NTD structure. The protein is shown as a cartoon model with the β-sheet I (β2/β3/β4), β-sheet II (β1/β5), and α-helices (α1 and α2) colored in dark green, light orange, and magenta, respectively. The thumb (residues I146–V158 in α2/β5) is colored in yellow.

We used the CSP as a function of DNA concentration to estimate the *K_d_* for the interaction with N-NTD (Table S8). In contrast to the nanomolar affinities measured in the absence of NaCl and inorganic phosphate (Table S7), we observed apparent dissociation constants in the order of micromolar, and yet, despite the difference in affinities the protein is active in all tested condition (Figure S2). The affinities for 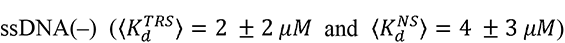 are higher than those for 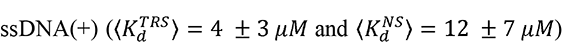, and ssTRSs display greater affinities than those observed for ssNSs. These observations agree with the fluorescence results.

### Hydrogen bonds and energy contributions to N-NTD:DNA binding

To better describe the interactions, we used the experimentally CSP-mapped binding interface to generate structural models of the N-NTD:DNA complexes using molecular docking followed by 2.0 μs molecular dynamics (MD) simulations. Structural models of the complexes generated by docking calculations correspond to the lowest energy structure from the cluster with the lowest HADDOCK score (fraction of common contacts > 0.8, interface-RMSD < 0.2 Å, and ligand-RMSD < 5.0 Å). Next, we performed 2 μs MD simulations for the docking models. This simulation time is long enough to probe the structural stability of the models, but not to explore all the conformational space. With that in mind, we showed that N-NTD complexes with both ssDNA and dsDNA reach stability at the end of the 2 μs MD simulations, exhibiting stable values of RMSD, number of protein-DNA contacts, and number of protein-DNA hydrogen bonds (Figure S7, S8, and S9). Thus, we determined representative structural models of the N-NTD:DNA complexes from the clustering of the MD trajectories (Figure 3G, 3H, and S10). The representative models are consensual in many aspects. For dsDNA, the major groove is recognized by the finger, and the 5’-end of the positive-sense strand is oriented toward the palm. We observed a difference of ∼25° in orientation of dsTRS and dsNS in the representative structure models (Figure 3I), positioning the dsNS close to the finger and dsTRS buried into the palm. The β-sheet II (β1/β5) interacts with the negative-sense DNA strand, especially with the TTT motif, suggesting that it might be involved in TRS recognition. For ssDNA, the 5’-end is oriented toward the palm and the finger interacts with a stretch of nucleotides spanning 5 residues from the 3^rd^ to the 7^th^ register. At least the last 3 nucleotides are free and do not interact with N-NTD. It is worth mentioning that the consensual structures of N-NTD:dsDNA complexes observed here are similar in ligand position and orientation compared to the consensual structure with the RNA duplex (PDB id 7ACS) (47).

The MD trajectories may report, not all, but important interactions related to the binding stability and specificity. Thus, we determined protein-DNA hydrogen bonds from the 2 μs MD trajectories and calculated their percentages of persistence along the simulations (Figure S8). Protein-DNA hydrogen bonds with persistence higher that 10% were counted with respect to amino acid and nucleotide residues (Figure 5, Table S1–S6). It is noteworthy that protein-DNA hydrogen bonds also report the presence of salt bridges between arginine or lysine residues and DNA phosphate groups (Figure 5G and 5H). For both dsDNA and ssDNA complexes, the presence of persistent protein-DNA hydrogen bonds mirrored the CSP results for the following N-NTD regions: the flexible N-terminal, the palm formed by β-sheet I, the finger, the β-sheet II, and the C-terminus. The remote region and the thumb displayed significant CSPs but no persistent protein-DNA hydrogen bonds or salt bridges (Figure 5). This feature reveals that non-polar interactions contribute to binding of the thumb to DNA and that the remote region is not directly involved in the recognition interface. Binding of DNA to the cleft formed between the finger and the palm is mostly mediated by positively charged residues. Residues R92, R107, Y172, and R177 are hydrogen bonded to the DNA oligonucleotide in most MD runs (Figure 5I). Remarkably, mutations R92E and R107E almost abolished N-NTD interaction with RNA (47). Residues E174 and K61 make hydrogen bonds almost exclusively with TRS. The negatively charged E174 is hydrogen bonded to the nitrogenous bases, positioned approximately in the middle of the two strands. By using molecular dynamic simulations, we suggested previously that this residue may contribute to the dsRNA melting activity of N-NTD (14).

**Figure 5.**
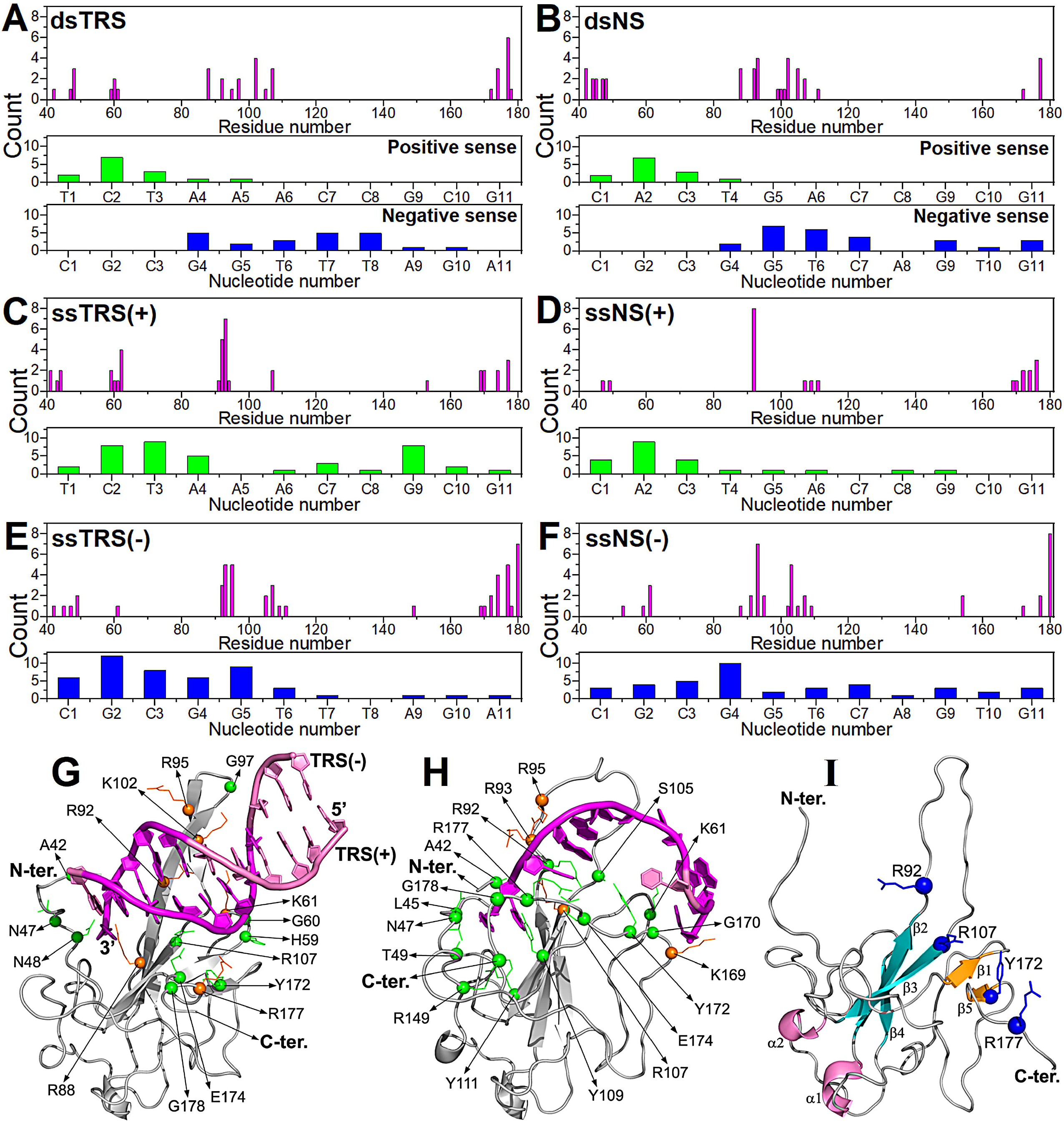
Protein-DNA hydrogens bonds involved in the structure models of the N-NTD:DNA complex. Count of protein-DNA hydrogen bonds with persistency higher that 10% were with respect to amino acid (magenta bars) and nucleotide residues (green and blue bars) for the interaction of the N-NTD with (A) dsTRS, (B) dsNS, (C) ssTRS(+), (D) ssNS(+), (E) ssTRS(–), and ssNS(–). The residues involved in the persistent protein-DNA hydrogen bonds (green sphere) and salt bridges (orange sphere) are indicated on the representative structural models of the (G) N-NTD:dsTRS and (H) N-NTD:ssTRS(–) complex. The protein is shown as a cartoon model in gray and DNA as a cartoon-ring model in light pink with TTT motif in magenta. (I) The most recurrent residues (R92, R107, Y172, and R177) involved in protein-DNA hydrogen bonds are indicated as blue spheres on the N-NTD structure. The protein is shown as a cartoon model with the β-sheet I (β2/β3/β4), β-sheet II (β1/β5), and α-helices (α1 and α2) colored in cyan, orange, and magenta, respectively.

The positive-sense strands of dsDNA (*middle*, Figure 5A and 5B) and ssDNA (Figure 5C and 5D) are mainly hydrogen bonded to N-NTD through their 5’ termini (nucleotide 1 to 4). Conversely, the negative-sense strands of dsDNA are mainly recognized by N-NTD through hydrogen bonds with their 3’ termini (nucleotides 4 to 9) (*bottom*, Figure 5A and 5B), which include the specific TTT motif for TRS. It is interesting to note that binding of N-NTD to positions 1 to 5 of the positive-sense strand and 4 to 9 of the negative-sense strand is maintained independently of the nucleotide sequence (TRS or NS), suggesting that the orientation of dsDNA with respect to the positively charged cleft is conserved and sequence independent. We observed consistently that the total number of protein-DNA persistent hydrogen bonds with ssDNA(–) (Figure 5E and 5F) is significantly larger than that with ssDNA(+) (Figure 5C and 5D). These results are in agreement with the observation that ssDNA(–) shows higher affinity to N-NTD and N-NTD-SR than ssDNA(+) (Table S7). The same tendency is observed for the negative-sense strands of dsDNAs (Figure 5A and 5B).

Due to the high content of charged residues and the above-mentioned electrostatic contribution (salt and phosphate dependence, Figure 2B and 2C) to DNA recognition, we decided to compute the theoretical Gibbs free energy of binding (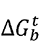) using Poisson-Boltzmann Surface Area (PBSA). This method enables to evaluate and discriminate the main protein residues and nucleotides that contribute to 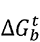. The most significant contributions come from the charged residues (Figure 6A to 6F) distributed throughout the protein (Figure 6G and 6H). Positively charged arginine and lysine residues contribute favorably, while negatively charged aspartate and glutamate residues contribute unfavorably. It is important to consider the unfavorable contributions because they may be responsible for the DNA/RNA duplex melting activity of N-NTD.

**Figure 6.**
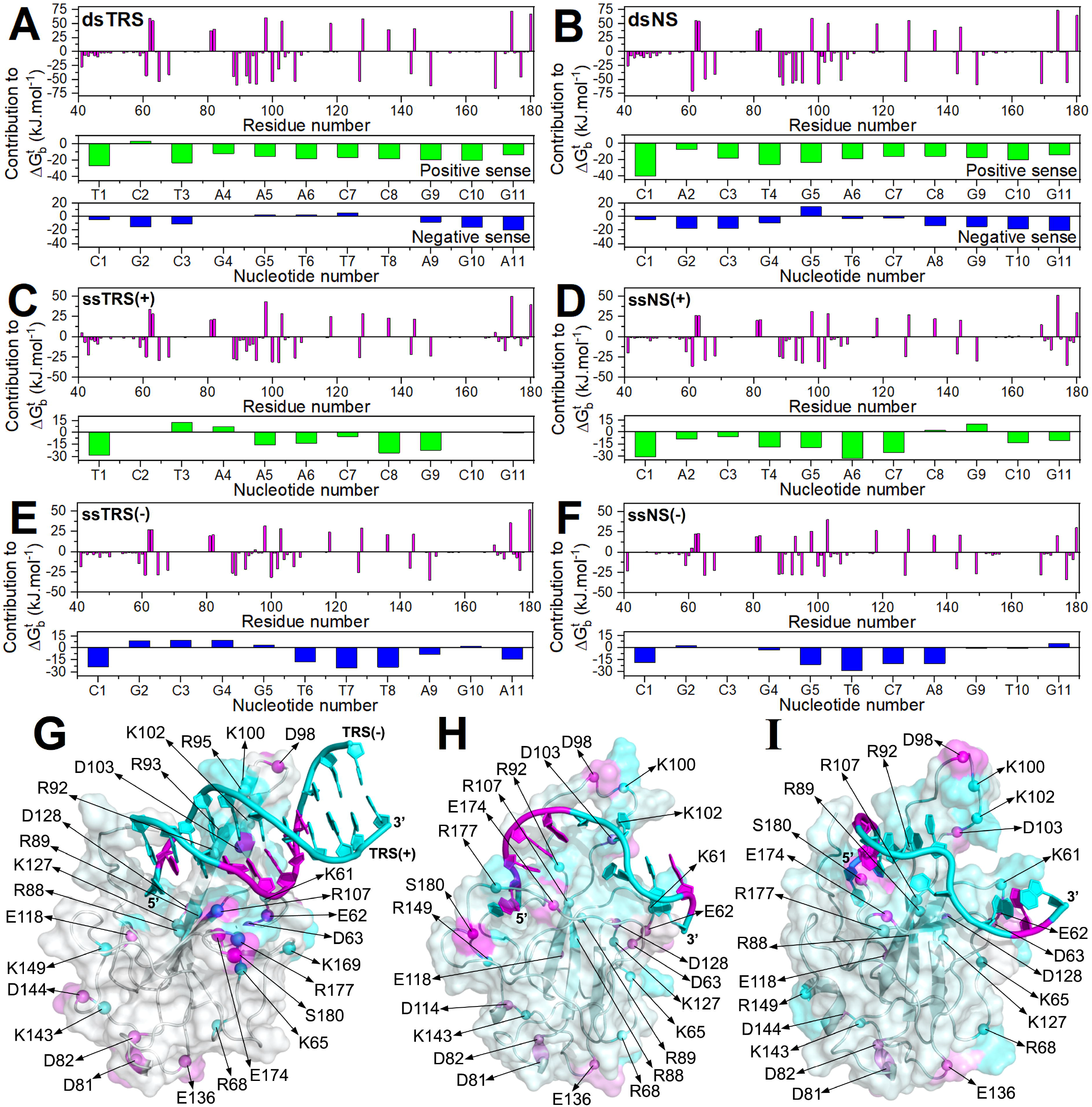
Energy contribution to Gibbs free energy change (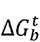) of N-NTD:DNA binding. Energy contribution of the amino acid (magenta bars) and nucleotide (green and blue bars) residues to 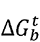 (in kJ·mol^−1^) for the interaction of the N-NTD with (A) dsTRS, (B) dsNS, (C) ssTRS(+), (D) ssNS(+), (E) ssTRS(–), and ssNS(–). Charged residues with most significant energy contribution to 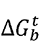 are indicated as spheres on the representative structural models of the complexes of N-NTD with (G) dsTRS, (H) ssTRS(–), and (I) ssTRS(+). The protein is shown as a cartoon model and translucid surface in gray with the favorable and unfavorable energy contribution of the residues colored in cyan and magenta gradient, respectively. The DNA molecule is displayed as a cartoon-ring model with the favorable and unfavorable energy contribution of the nucleotides colored in cyan and magenta, respectively.

It is also important to analyze the contributions to 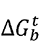 coming from the nucleic acid. For dsDNA, the contribution is mostly favorable for the positive-sense strand. For the negative-sense strand, the contribution is favorable at the 5’ and 3’ termini, but for positions 4 to 8 it varies according to the nucleotide sequence, being less favorable or near-zero for dsNS and unfavorable for dsTRS. For dsTRS, the stretch of nucleotides at positions 4 to 8, which contains the specific TTT motif, is hydrogen bonded to the β-sheet II (β1/β5) (Figure 5A and 5G). The same is not observed for dsNS, because it is tilted ∼25° away from β-sheet II (β1/β5) (Figure 3I). In this protein region, we observed charged residues that contribute unfavorably to 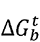, such as E174, which was identified in our previous study with dsRNA (14).

For ssDNAs, both positive and negative-sense strands contributed similarly to 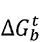, being slightly more favorable for the positive-sense strands. Interacting as a duplex, the positive-sense strands (TRS and NS) showed favorable contribution all along its sequence (Figure 6A and 6B); however, when interacting as a ssDNA, the contribution is less favorable (Figure 6C and 6D). For the negative-sense strands (TRS and NS), the nucleotides at positions 4 to 8 contribute unfavorably when interacting as a duplex DNA (Figure 6A and 6B), while favorably as a single strand (Figure 6E and 6F). This is remarkable and agrees with the hypothesis that these unfavorable contributions (TTT motif and β-sheet II) may play a key role in N-NTD melting activity. The contrasting behavior for each strand in the single or double-stranded DNA may be attributed to the fact that the dsDNA has a better-defined positioning when compared to the ssDNA, due to the presence of secondary structure in the duplex. The difference of ∼25° in orientation of dsTRS and dsNS (Figure 3I) may serve as an indicative of specificity of the TRS, contributing to the positioning β-sheet II, E174 for instance, close to TTT motif. This difference in positioning seems to be more relevant for the specificity than the difference in binding affinities between dsTRS and dsNS.

### N-NTD-SR phase separation is triggered by the SR motif and modulated by TRS binding

We investigated the ability of N-NTD and N-NTD-SR to form biomolecular condensates by liquid-liquid phase separation (LLPS) as a consequence of their interaction with specific or non-specific nucleic acids. Indeed, full-length N has been shown to form liquid droplets induced by RNA interaction (20–23). Interestingly, N from several coronaviruses is predicted to undergo LLPS by the catGRANULE algorithm (Figure S11A), while N-NTD-SR exhibits the highest LLPS propensity, most strikingly the SR-rich motif (Figure S11B). These characteristics prompted us to investigate whether the N-terminal domain contributes to the nucleic acid-driven LLPS of N, the role played by the SR-rich in phase separation, and the influence of nucleic acid binding specificity and stoichiometry on LLPS modulation. Notably, in the simultaneous presence of 10% PEG-4000 and a long non-specific RNA, we observed numerous spherical micron-sized condensates for N-NTD-SR only (Figure S12B). The same was not observed for N-NTD (Figure S12A), suggesting that the SR-rich region is important for inducing RNA-driven LLPS under macromolecular crowding. Neither N-NTD nor N-NTD-SR alone formed condensates in aqueous buffer (20 mM Tris-HCl pH 7.5, 30 mM NaCl) even with 10% PEG-4000. We also did not observe condensates in the presence of the long non-specific RNA without the crowding agent (Figure S12). The dependence of PEG-4000 for N-NTD-SR:RNA condensation was confirmed quantitatively by turbidity measurements (Figure S12C and S12D). To further dissect the impact of RNA concentration in promoting N-NTD-SR LLPS, we quantified the number of condensates per 100 µm^2^ area and their respective sizes (in µm^2^) at five protein:RNA molar ratios. We observed a progressive increase in the number of liquid droplets visible by microscopy accompanied by an increase in turbidity upon the increase in RNA concentration, starting at 2:1 N-NTD-SR:RNA molar ratio (Figure S13A and S13C). Specifically, an excess of RNA (1:2 N-NTD-SR:RNA) resulted in 305 ± 7 condensates in a 100 µm^2^ area (*top*, Figure S13B). This number was only 1.5 times higher than that observed at critical concentration for RNA-driven N-NTD-SR LLPS, i.e. at the 2:1 N-NTD-SR:RNA stoichiometry. The 1:2 N-NTD-SR:RNA ratio showed the highest dispersion on condensates size, consonant to large condensates formed by fusion (Figure S13B, bottom graph).

Since nucleic acid structure and sequence can finely tune the phase behavior, we followed this phenomenon in the presence of ssTRS and dsTRS DNAs (Figure 7). dsTRS (Figure 7A and Figure 7D) triggered significant N-NTD-SR LLPS at equimolar concentration (1:1 N-NTD-SR:dsTRS), whereas at 1:2 molar ratio, the condensates dissolved, a characteristic behavior of reentrant liquid condensation as reported for RNA-binding proteins containing disordered regions rich in arginine (48, 49). The appearance of condensates is coupled with an increase in turbidity (after 30 minutes incubation) at 1:2 stoichiometry, followed by a steep decrease in the presence of molar excess of DNA, confirming the microscopy data (Figure 7E). We also investigated the effect of the single-stranded counterparts of the duplex TRS. Incubation of N-NTD-SR with either ssTRS(+) (Figure 7B) or ssTRS(–) (Figure 7C) only resulted in the appearance of numerous condensates observed by microscopy as well as a significant increase in turbidity at 1:2 protein:DNA stoichiometry (Figure 7E). Additionally, condensates formed by N-NTD-SR:dsTRS displayed a smaller and more homogenous size as evidenced by their narrower size distribution in comparison to the ssTRSs (at 1:1 with dsTRS 5.4 ± 4.4 µm^2^; at 1:2 with ssTRS(+) 11 ± 12.1 µm^2^ and 1:2 with ssTRS(–) 10.4 ± 8.2 µm^2^). To confirm that the base-paired dsTRS was demixed together with N-NTD-SR, we used the probe DAPI, which shows about 20 times higher fluorescence quantum yield when bound to dsDNA (50). Indeed, just N-NTD-SR:dsTRS samples showed the extrinsic blue fluorescence emission (Figure 7D, insets). Fluorescence micrographs acquired with the same settings were obtained for N-NTD-SR in the presence of the long non-specific RNA (Figure S13B, insets) or with addition of ssTRS(+) or ssTRS(–) (Figure 7D, insets) and did not show DAPI fluorescence. To unveil the nature of the interactions that drive N-NTD-SR:dsTRS condensation at equimolar stoichiometry, we changed the solution composition (Figure S14). The addition of a high concentration of 1,6-hexanediol (10% (w/v) HD) decreased the number of droplets by about 30% (203.2 ± 8.2 droplets in absence of HD *versus* 146.8 ± 12.1 droplets with 10% HD) (Figure S14). This aliphatic alcohol is known to disrupt the weak hydrophobic contacts involved in the liquid condensates, whereas solid-like aggregates (51) and lipid membranes (52) are resistant to HD treatment. Proteins such as hBex3 (44), PrP^90-231^ (43), and RNA polymerase (53), to name a few scaffolds of LLPS, have phase separation abrogated by 10% HD, thus, electrostatic contacts should play a dominant role for N-NTD-SR:DNA LLPS. Indeed, upon incubation with 300 mM NaCl, N-NTD-SR:dsTRS LLPS was totally disassembled (Figure S14A). Interestingly, in buffer containing 20 mM sodium acetate at pH 5.5, droplet formation was higher, and condensates had marked circular morphology (Figure S14). Specifically, the droplet number increased by 32% compared to pH 7.5 (203.2 ± 8.2 droplets at pH 7.5 *versus* 298.6 ± 16.8 droplets at pH 5.5), suggesting that acidic pH induces nucleic acid-driven N-NTD-SR LLPS.

**Figure 7.**
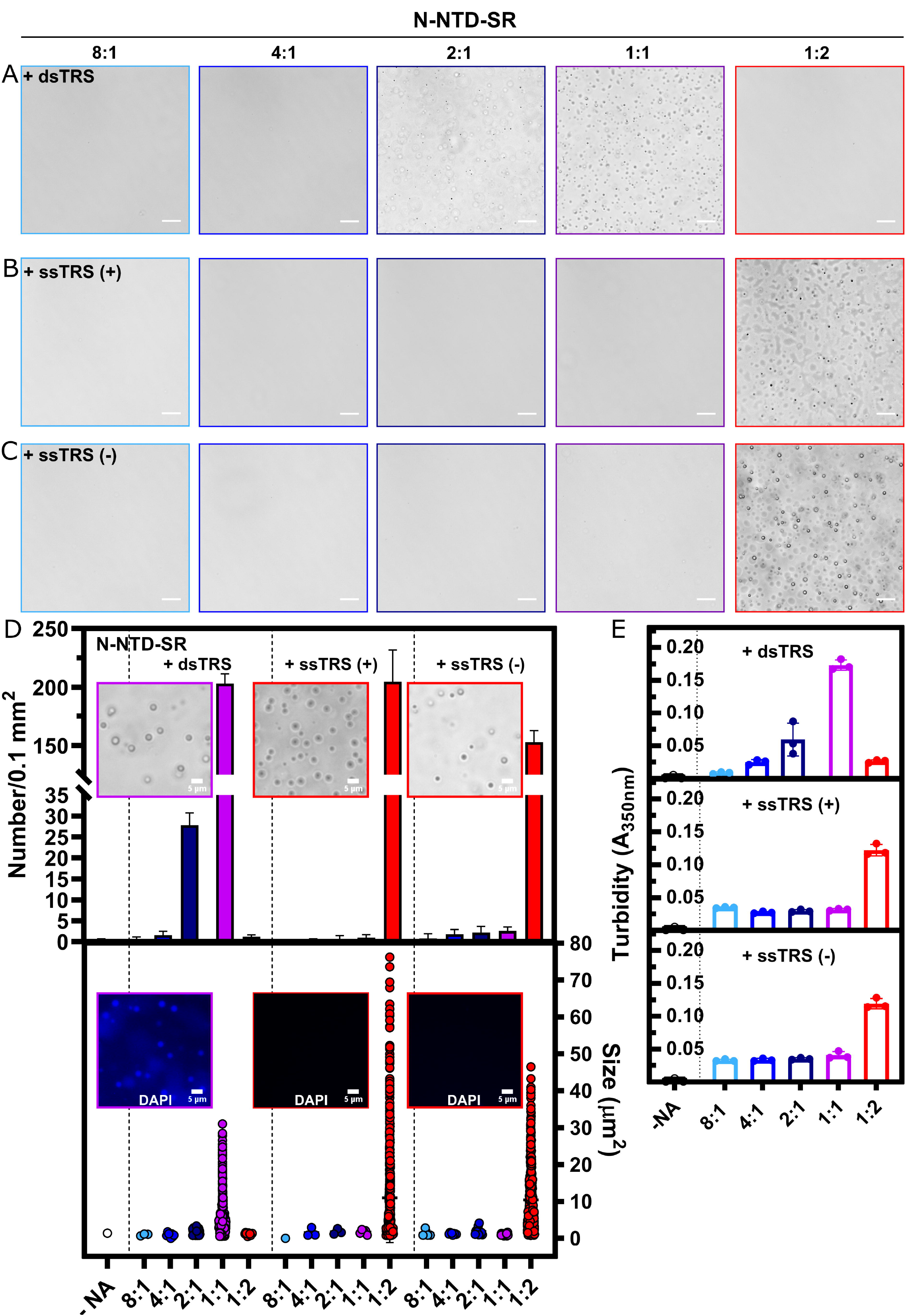
The high-affinity duplex TRS triggers N-NTD-SR phase separation at a lower protein:DNA stoichiometry. Representative phase contrast micrographs of 20 µM N-NTD-SR in presence of dsTRS (A); ssTRS(+) (B) or ssTRS(–) (C) at the following protein:DNA stoichiometries: 8:1 (light blue); 4:1 (marine blue); 2:1 (dark blue); 1:1 (purple); 1:2 (red), respectively, in 20 mM Tris-HCl buffer (pH 7.5) with 30 mM NaCl. (D) Top graph: Mean number of condensates ± S.D. per 100 µm^2^ area (n= 5 images). Top insets: Phase contrast microscopy in the presence of DAPI for conditions with the highest number of liquid droplets (1:1 for the dsTRS; 1:2 for ssTRS(+), and 1:2 for ssTRS(–)). Bottom graph: Scatter plot from the size of condensates (represented as area in µm^2^) obtained from micrographs analysis. The following number of condensates were measured: -NA (1); for dsTRS at 8:1 (N= 3); 4:1 (N= 8); 2:1 (N= 132); 1:1 (N=1014) and 1:2 (N= 11); for ssTRS(+) at 8:1 (N= 1); 4:1 (N= 3); 2:1 (N= 3); 1:1 (N= 5) and 1:2 (N= 1019); for ssTRS(–) at 8:1 (N= 4); 4:1 (N= 9); 2:1 (N= 11); 1:1 (N= 21) and 1:2 (N= 759). Bottom inset: corresponding DAPI emission from the top graph insets images (DAPI stains condensates in presence of dsTRS and no fluorescence were observed for the ssTRSs). All conditions contained 10% (w/v) PEG-4000. Scale bar, 20 µm apart from insets (5 µm). (E) Phase separation of 20 µM N-NTD-SR in 20 mM Tris-HCl buffer (pH 7.5) with 30 mM NaCl in the presence of 10% (w/v) PEG-4000 monitored by absorbance measurements at 350 nm as a function of increasing concentrations of the specific DNA oligonucleotides dsTRS (top graph), ssTRS(+) (middle graph), and ssTRS(–) (bottom graph). The protein:DNA stoichiometries are the same as in (A).

To investigate the role of sequence specificity in promoting the N-NTD-SR LLPS, we followed the condensation process using the NS DNAs (Figure S15). In agreement with dsTRS, the relevant condition for LLPS was at 1:1 protein:DNA stoichiometry. In addition, excess of dsNS dissolved the condensates (127.2 ± 13.9 condensates/µm^2^ at 1:1 *versus* 18.4 ± 3.6 condensates/µm^2^ at 1:2). This behavior was confirmed by turbidity measurements (Figure S15E). However, condensates were not homogeneously spherical, as the ones formed with specific dsTRS, and most of them wetted the coverslip surface. Curiously, when observing the entire cover of the glass slide from 1:1 N-NTD-SR:dsNS samples, we observed a few crystals. Since all images were obtained after 30 minutes incubation, we sought to understand whether crystal formation would be enhanced with prolonged incubation. After 2 hours of incubation, we observed crystals presenting DAPI staining (Figure S15D, inset). Condensates are supersaturated and estimated to be 10 to 300 times enriched in macromolecules compared to the diffuse phase (54), thus crystallization is thermodynamically favored. The buffer with addition of 10% PEG-4000 did not show any artifact (Figure S12B, top right image), as well as isolated oligonucleotides used in the experiments were excluded to give rise to condensates as shown for dsNS (Figure S12B, bottom right). The N-NTD-SR:dsNS transition from liquid-like condensates to crystals suggests that the structural properties of the separated phase are different from the N-NTD-SR:dsTRS, once again highlighting the specificity of interaction. Since both dsTRS and dsNS have secondary structure, the higher affinity of N-NTD-SR toward dsTRS might play a role in maintaining the liquidity and consequent dynamics of the protein:nucleic acid condensates. Additionally, we did not observe either condensate formation or increase in turbidity in the presence of the ssNS(+) or ssNS(–) at the five different molar ratios examined (Figures S15B, S15C, and S15E). Therefore, an indicative of sequence specificity may play a pivotal role for DNA-driven N-NTD-SR LLPS since only ssTRS(+) and ssTRS(–) were able to trigger LLPS and not the ssNSs.

## DISCUSSION

The nucleocapsid N protein is well characterized for its interaction with RNA, which is essential to understand two important biological processes for the viral cycle: (i) the assembly of the helical ribonucleoprotein (RNP) complex, and (ii) the discontinuous transcriptional mechanism. N’s dsRNA melting activity enables template switch during discontinuous transcription (7, 10). In addition, its ability to phase separate creates a membraneless compartment (liquid-like condensates) that regulates transcription and replication. Previous studies have shown that N binds to single and double-stranded DNA as RNA mimetics (55, 56). Takeda and cols. (2008) and Zhou and cols. (2020) characterized the DNA interaction with the C-terminal domain of N (N-CTD) from SARS-CoV (57) and SARS-CoV-2 (58), respectively. Here, we describe at molecular detail the binding and activity (melting and formation of liquid condensates) of N-NTD and N-NTD-SR toward specific and non-specific DNAs.

Structural and binding studies revealed that solvent-exposed charged residues and electrostatic interactions are the main driving forces for N:nucleic acid complex formation (10, 47, 55, 57, 58). In line with that, we observed that formation of the N-NTD:DNA complex is NaCl and inorganic phosphate dependent (Figure 2B and 2C). In addition, dissociation constant increases from nanomolar in 20 mM Bis-Tris buffer (no salt and pH 6.5, Table S7) to the micromolar range in 20 mM sodium phosphate buffer (pH 6.5, Table S8) containing 50 mM NaCl, while maintaining melting activity (Figure S2). This is in agreements with kinetic simulations of the melting activity that suggests the activity does not depends on the binding affinity so long *K_d_* < 10^−1^ M^−1^ (14). It is worth mentioning that the low micromolar *K_d_* values are similar to those described for RNA binding to N-NTD of SARS coronavirus (47, 59) and Mouse Hepatitis Virus (MHV) (7), a virus closely related to SARS-CoVs. The MD simulations, based on the experimental binding data obtained by NMR, revealed that electrostatic interactions constitute the main energetic contribution for N-NTD:DNA complex stabilization, which happens through the formation of hydrogen bonds between opposite charged residues, *i.e.* salt bridges. Despite the positive electrostatic potential on the binding interface (14), MD simulations also showed the importance of unfavorable contributions from negatively charged residues to 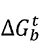. Accordingly, Dinesh and cols. (2020) showed that R92E and R107E mutations, located at the binding cleft, lead to a decrease in N-NTD-RNA binding affinity, while E174R promotes an increase (15). We also observed from PBSA analysis that short and long-range contributions of charged residues to 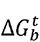 play a key role in protein-DNA binding affinity and perhaps in dsDNA melting activity.

We provide the first evidence that N-NTD and N-NTD-SR are active upon binding to DNA. Both constructs show dsDNA melting activity, and N-NTD-SR forms liquid condensates in the presence of DNA in a crowded physiological buffer. It is noteworthy that these activities were reported for RNA only (7, 10, 20, 21, 60), and that the ability to melt dsRNA and to form condensates in the presence of RNA are important biological functions of N to spatiotemporally regulate the transcription and replication machinery. We hypothesize that the similar positioning of DNA and RNA in the binding cleft (finger/palm) is the determining factor for the activities of N-NTD and N-NTD-SR, making the interaction with DNA also a trigger for LLPS. This proposal is based on the similarity between the CSP profiles obtained for N-NTD/N-NTD-SR when interacting with DNAs (Figure 3 and S5) and RNAs (15), both single and double-stranded. As a consequence, the structural models for N-NTD:DNA (Figures 3G, 3H, and S10) and N-NTD:RNA complexes (15) are remarkably similar in position and orientation of both single and double-stranded nucleic acids. We speculate that the conformational space explored by the single-stranded nucleic acid dynamics might compensate for the occupancy of the double helix in the binding cleft. Taking this interpretation together with the intrinsic protein dynamics, especially at the binding site where a tweezer-like motion between the finger and α2/β5 loop takes place (14), we speculate that the entropically driven ssTRS interaction (Figure 2D) may be due to the conformational freedom of both protein and ssDNAs. In contrast, the better-defined position and orientation of dsTRS in its enthalpically driven interaction (Figure 2D) would have the protein conformational dynamics as major contribution since DNA duplex destabilization is an entropically driven process. The PCA analysis of the MD simulations showed that the conformational space probed by the ssTRS-bound N-NTD is significantly higher than by the dsTRS-bound N-NTD (Figure S16). Our results suggest that the dsDNA defined positioning impacts both the formation of liquid droplets and the melting activity. For LLPS, the presence of nucleic acid secondary structure (dsDNA/dsRNA) favors adequate positioning and perhaps this is linked to the fact that dsTRS induces LLPS at lower DNA concentration when compared to ssTRS (2:1 for N-NTD-SR:dsTRS and 1:2 for N-NTD-SR:ssTRSs). For the melting activity, the defined positioning is also important. We observed dsDNA melting activity at low and high salt and phosphate concentrations (Figure 1 and S2), for which we showed changes in affinity ranging from nano- to micromolar, suggesting that it is not the difference in affinity that leads to the melting activity, but a property of the interaction itself. Protein dynamics and the defined positioning of dsDNA/dsRNA may be the key for this N protein biological function.

Unlike homeodomains found in eukaryotic transcription factors that possess well-characterized sequence-specific DNA binding (61), CoV N and its CTD are described as non-specific nucleic acid-binding proteins as they bind to RNA and DNA, both single and double-stranded (10, 55, 59). This characteristic corroborates the multifunctional role played by N during the viral cycle (11). Our results for N-NTD/N-NTD-SR binding to DNA reinforce the importance of non-specific charged interactions, such as those observed for RNA (7, 10, 62). In general, we noticed modest differences comparing *K_d_* values for TRS and NS DNAs, as well as for ssDNAs and dsDNA, which might suggest a sequence-specific recognition of TRS with respect to NS and an affinity preference for ssDNA(–) over ssDNA(+) and dsDNA (Table S7 and S8). However, considering the difference between specific and non-specific dsDNA binding for homeodomains, there are a ratio of hundreds of times of *K_d_* values (63) that defeats our suggestion of sequence-specific recognition by N-NTD or N-NTD-SR. According to these observations, we cannot explain the specificity based on differences in the affinities, thus we propose the existence of the following indicatives of sequence-specific recognition for TRS: (i) the unique encounter of unfavorable energetic contributions to 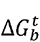 along with protein-dsTRS hydrogen bonds between the TTT motif and β-sheet II, notably for E174 that might trigger dsRNA/dsDNA melting; (ii) a well-defined pattern for the thermodynamic profile (ΔH and ΔS > 0 for ssTRSs, and ΔH and ΔS < 0 for dsTRS, Figure 2D) for TRS interaction with N-NTD/N-NTD-SR; and (iii) the preference for TRS over NS DNAs in the formation of liquid condensates. We observed LLPS for ssTRSs but not for ssNS, which might be explained by the difference in affinity to ssNSs (Table S7 and S8), and crystalline aggregates for dsNS, despite the occurrence of LLPS for both dsTRS and dsNS.

In conclusion, our results contribute to a detailed understanding of the molecular characteristics of binding and activities of the SARS-CoV-2 N protein toward nucleic acids. We described the interaction of N-NTD/N-NTD-SR with single and double-stranded DNA, as well as its dsDNA melting activity and DNA-induced LLPS. In agreement with RNA studies, we characterized the importance of electrostatic interactions for N-NTD(-SR):DNA complex formation and stabilization. Our study also shed light on indicatives of specificity in a protein that is widely described as nonspecific. We suggest that interaction of dsDNA/dsRNA in the binding cleft presents a defined position and orientation of the double helix. For TRS, the melting activity of N-NTD/N-NTD-SR does not seem to be related to the differences in binding affinity to DNA (or RNA), but to differences in positioning of the TTT(UUU) motif in the duplex close to β-sheet II (Figure 3I), where E174 (conserved in all human CoVs, Figure S17) might act as a trigger for duplex destabilization. The activity toward DNA may serve as a starting point for the design of inhibitors and/or aptamers against a therapeutic target essential for the virus infectivity (7). In addition, it opens a new avenue of investigation arguing if DNA is indeed a cellular target important for the virus cycle. It is worth noting that N protein of SARS-CoV and SARS-CoV-2 contains conserved nuclear localization signals (NLS) and nuclear export signal (NES) (Figure S17) (64), despite the fact that SARS-CoV N has not been observed inside the nucleus (65, 66).

## ACKNOWLEDGMENTS

The author IPC gratefully acknowledges the financial support by postdoctoral fellowship from FAPERJ and the PROPe UNESP. The authors are grateful for the access to the Santos Dumont supercomputer at the National Laboratory of Scientific Computing (LNCC), Brazil. We also acknowledge the Covid-19 NMR Consortium (https://covid19-nmr.de/) for providing an excellent environment for scientific discussions. This work is a collaborative effort of the Rio BioNMR Network (https://www.rmnrio.org/).

