## Supplementary material for "Structure insights, thermodynamic profiles, dsDNA melting activity, and liquid-liquid phase separation of the SARS-CoV-2 nucleocapsid N-terminal domain binding to DNA"

***To whom correspondence should be addressed:**

### SUPPLEMENTARY MATERIAL

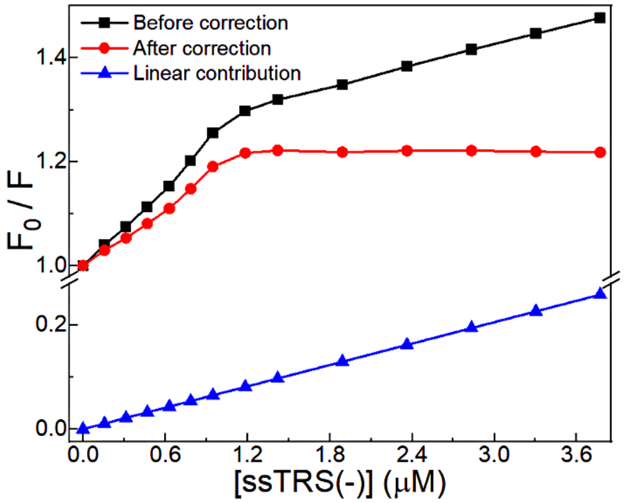


**Figure S1.** Removal of the upward linear contribution (blue triangle) from the fluorescence titration curves (black square) resulting in the corrected binding isotherm (red circle).

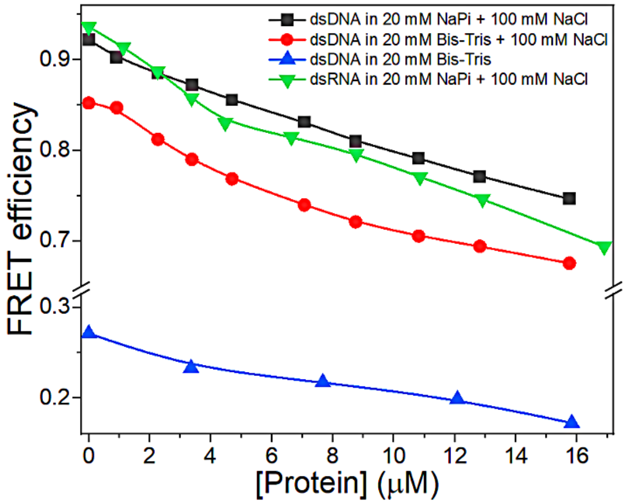


**Figure S2.** FRET efficiency as a function of the total concentration of N-NTD-SR in 20 mM sodium phosphate buffer (pH 6.5) containing 100 mM NaCl for dsDNA TRS (black squares), 20 mM Bis-Tris (pH 6.5) containing 100 mM NaCl for dsDNA TRS (red circles), 20 mM sodium phosphate buffer (pH 6.5) containing 100 mM NaCl for dsRNA TRS (blue triangle), and 20 mM Bis-Tris (pH 6.5) for dsDNA TRS (green inverted triange).

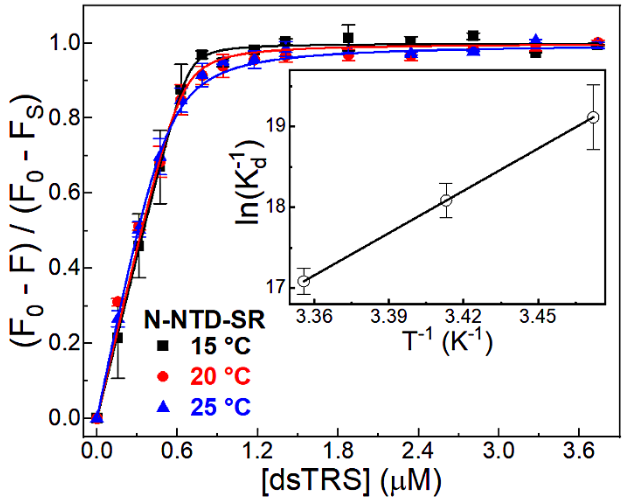


**Figure S3.** Fluorescence quenching changes of N-NTD as a function of the dsTRS concentration in 20 mM Bis-Tris buffer (pH 6.5) at 15, 20, and 25 °C. Each point on the binding isotherm represents the average and standard error calculated from duplicate measurements. The continuous lines denote the theoretical curves globally adjusted to the experimental data. The inset shows the van't Hoff plot determined the enthalpy change value for the N-NTD/ssTRS(–) complex.

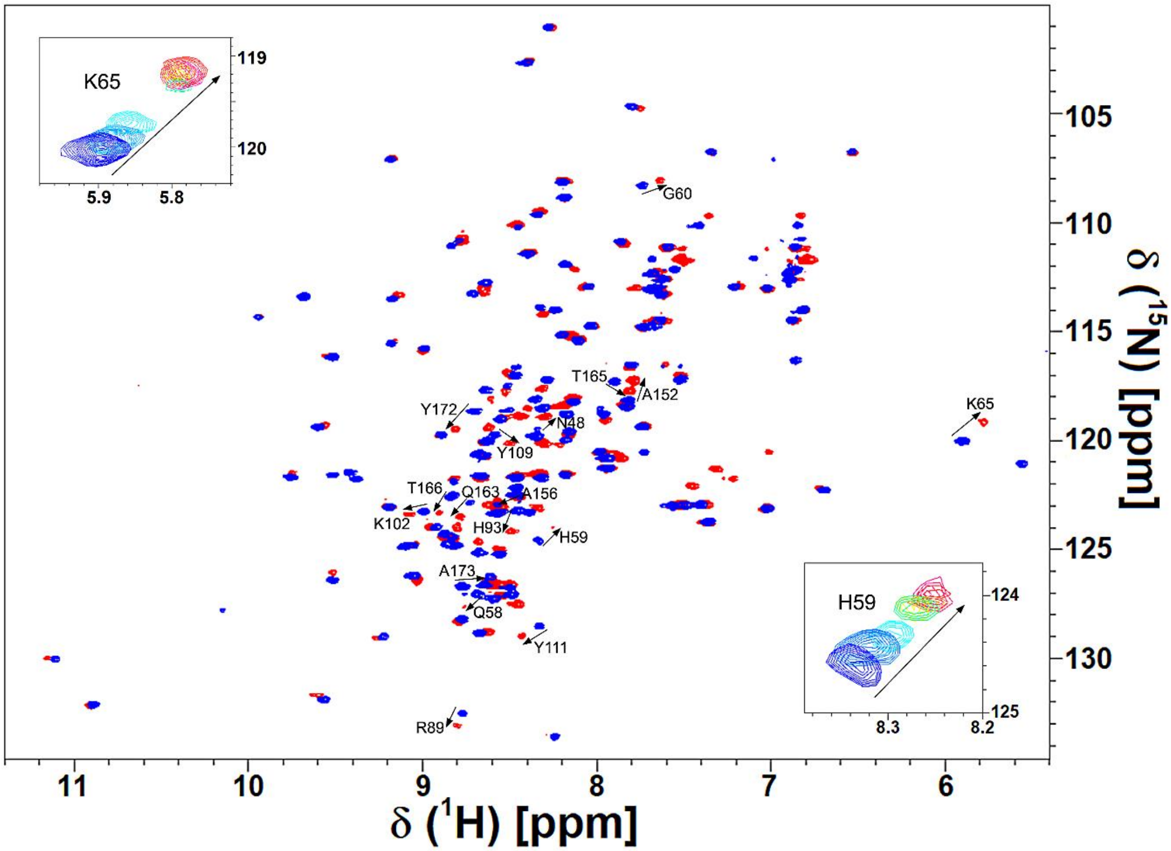


**Figure S4.** Two-dimensional [$^{1}$H-$^{15}$N] HSQC spectra of the free (blue) and ssTRS(–)-bound (red) [U-$^{15}$N]N-NTD collected by using an NMR spectrometer operating at $^{1}$H frequency of

600.033 MHz at 20°C. The arrows indicate the residues that presented a chemical shift perturbation (CSP) upon DNA binding higher than $\Delta\delta_{ave}$ + SD. (Insets) Titration effect on the behavior of the exchange regime for H59 and K65 upon ssTRS(–) binding. The spectra were recorded at a protein concentration of 70 μM (dark blue) and at an DNA concentration of 9.8 μM (light blue), 23.8 μM (cyan), 53.9 μM (green), 70 μM (yellow), 109.9 μM (orange), 174.4 μM (pink), and 209.3 μM (red).

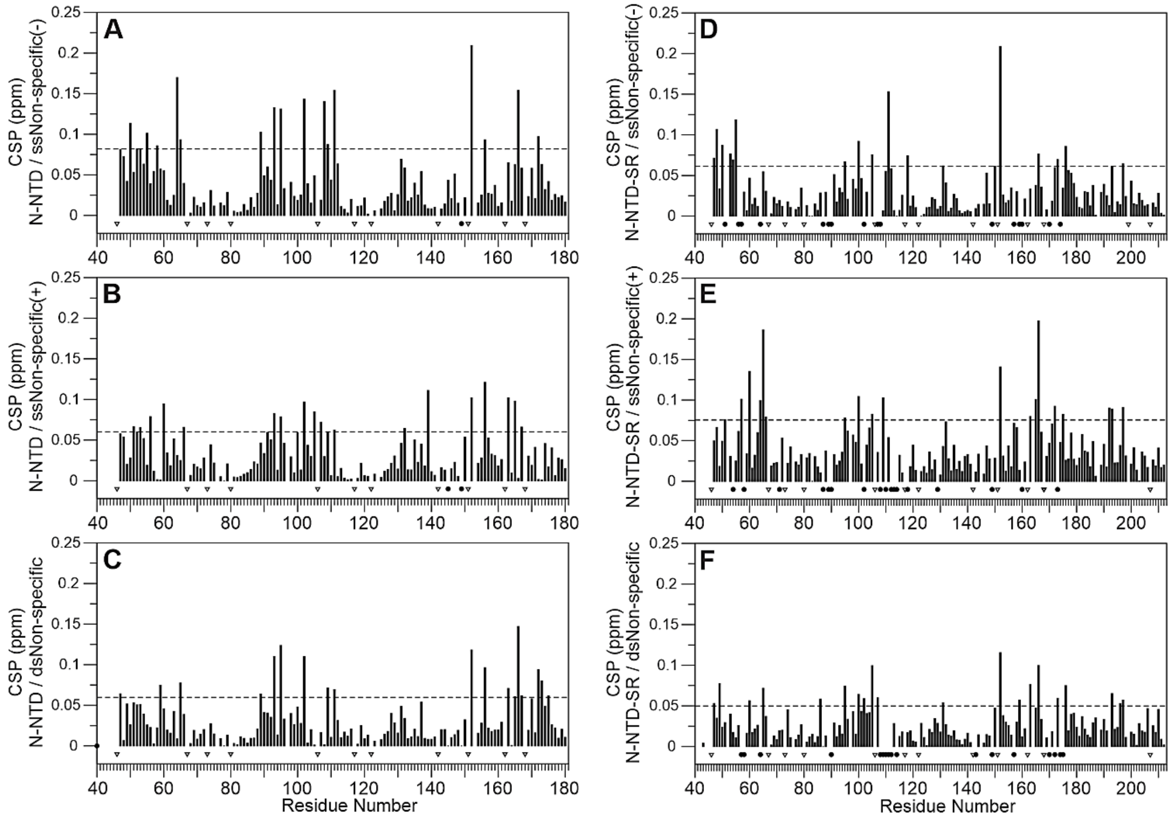


**Figure S5.** Chemical Shift Perturbation (CSP) of N-NTD and N-NTD-SR protein upon interaction with non-specific (NS) DNA. Chemical Shift Perturbation (CSP) of N-NTD (A, B, and C) and N-NTD-SR (D, E, and F) upon interaction with ssNS(–) (A, D), ssNS(+) (B, E), and dsNS (C, F). A chemical shift perturbation higher than the average CSP value ($\Delta\delta_{ave}$) plus one standard deviation (SD) (dotted line) was considered statistically significant and used as experimental constraints in subsequent docking calculations. The proline residues (46, 67, 73, 80, 106, 117, 122, 142, 151, 162, 168, 199 and, 207) are indicated by triangle-down. Resonances signals broadened beyond detection upon NS titration are represented by the filled black circles.

| protein-DNA complex/sequence | N | N | T | A | S | F | T | A | L | T | Q | H | G | K | L | K | F | Y | R | A | T | R | R | I | R | G | K | K | D | L | S | R | W | Y | Y |
|---|---|---|---|---|---|---|---|---|---|---|---|---|---|---|---|---|---|---|---|---|---|---|---|---|---|---|---|---|---|---|---|---|---|---|---|
| >N-NTD+ssTRS(-) | | 48 | | | | | | | | # | 58 | 59 | 60 | | # | 65 | | | 89 | | | | 93 | | | | | 102 | | | | | | 109 | 111 |
| >N-NTD+ssTRS(+) | | | | | 51 | 53 | 54 | | 56 | # | # | # | | 61 | # | 65 | | | 89 | | | 92 | 93 | 94 | | | | | | | | | | 109 | 111 |
| >N-NTD+dsTRS | | | | | 51 | | | | 56 | 57 | | 59 | | | 64 | 65 | | | | # | | 92 | | | 95 | | | | | | | | 108 | | |
| consensus | | | | | : | | | | : | * | : | * | | | * | * | | | : | | | : | : | | | | | | | | | | | : | : |
| >N-NTD+10mer-ssTRS(+)(RNA)§ | | | | | | | | | 56 | | | | 60 | 61 | | 65 | 66 | | | 90 | | | 93 | 94 | 95 | | 100 | 102 | 103 | 104 | | | | | |
| >N-NTD+7mer-ssTRS(+)(RNA)§ | | | | | | | | | 56 | 57 | | | 60 | | | 65 | 66 | | | 90 | | | 93 | 94 | 95 | | | 102 | 103 | 104 | | | | | |
| >N-NTD+7mer-dsNS(RNA)§ | | | | 50 | | | | | 56 | 57 | | | | | | 65 | 66 | | | | | 92 | 93 | 94 | | 96 | | | 103 | | 105 | 107 | 108 | | |
| >N-NTD+ssNS(-) | 47 | | | 50 | | 53 | | 55 | | | 58 | | | | 64 | 65 | | | 89 | | | | 93 | | 95 | | | 102 | | | | | 108 | 109 | 111 |
| >N-NTD+ssNS(+) | 47 | | | | 51 | 53 | | | 56 | | | | 60 | | | | 66 | | | | | | 93 | | 95 | | 100 | 102 | | | 105 | 107 | | 109 | 111 |
| >N-NTD+dsNS | 47 | | | | | | | | | | | 59 | | | | 65 | | | 89 | | | | 93 | | 95 | | | 102 | | | | | | 109 | 111 |
| consensus | * | | | | | : | | | | | | | | | | : | | | : | | | | * | | * | | | * | | | | | | * | * |
| >N-NTD-SR+TRS(-) | | 48 | | | | | 54 | | | | | | | | 64 | 65 | | | | 90 | 91 | | 93 | | | | 100 | 102 | | | | | | | 111 |
| >N-NTD-SR+TRS(+) | | | | | | 53 | 54 | | 56 | 57 | | | | | | 65 | | | | | | 92 | | | 95 | | | 102 | | | | | | 109 | 111 |
| >N-NTD-SR+dsTRS | | | | | | | | | | | | | 60 | | 64 | 65 | | | 89 | 90 | | | | | 95 | | 100 | 102 | | | | | 108 | 109 | 111 |
| consensus | | | | | | | : | | | | | | | | : | * | | | | : | | | | | : | | : | * | | | | | | : | * |
| >N-NTD-SR+ssNS(-) | 47 | 48 | | 50 | # | 53 | 54 | 55 | # | | | | | | # | | | | | # | | | | | 95 | | 100 | | | | 105 | | | | 111 |
| >N-NTD-SR+ssNS(+) | 47 | 48 | | 50 | | 53 | 54 | 55 | | | | | | | | 65 | | | | # | | | | | 95 | | 100 | | | | 105 | | | | 111 |
| >N-NTD-SR+dsNS | 47 | | 49 | | # | | # | # | # | # | # | | 60 | # | # | 65 | | 86 | | # | | | | | 95 | | 100 | 102 | | | 105 | 107 | # | # | # |
| consensus | * | : | | : | : | : | * | * | : | | | | | | : | : | | | | : | | | | | * | | * | | | | * | | | | * |

| protein-DNA complex/sequence | Y | G | E | I | W | A | L | R | N | A | A | I | L | Q | Q | T | T | L | G | Y | A | E | G | S | R | R | S | S |
|---|---|---|---|---|---|---|---|---|---|---|---|---|---|---|---|---|---|---|---|---|---|---|---|---|---|---|---|---|
| >N-NTD+ssTRS(-) | | | | | | | | | | 152 | 156 | | | | 163 | 165 | 166 | | | 172 | 173 | | | | | | | |
| >N-NTD+ssTRS(+) | | | | | | 134 | | | | 152 | 156 | | | | 163 | 165 | 166 | | | # | 173 | # | | 176 | | | | |
| >N-NTD+dsTRS | | 114 | | | | | | | 150 | 152 | 156 | | | | 163 | 165 | 166 | | | 172 | 173 | | | | 177 | | | |
| consensus | | | | | | | | | | * | * | | | | * | * | * | | | * | * | | | | | | | |
| >N-NTD+10mer-ssTRS(+)(RNA)§ | | | | | | | | 149 | | | 156 | 157 | | 160 | | 165 | 166 | 167 | | | | | 175 | | 177 | | | |
| >N-NTD+7mer-ssTRS(+)(RNA)§ | | | | | | | | 149 | | 152 | | 157 | | | | 165 | 166 | 167 | | | | | | | | | | |
| >N-NTD+7mer-dsNS(RNA)§ | | | | | | | | 149 | 150 | 152 | | | | | 163 | 165 | 166 | 167 | | 172 | 173 | | | | | | | |
| >N-NTD+ssNS(-) | | | | | | | | | | 152 | 156 | | | | | | 166 | | | 172 | | | | | | | | |
| >N-NTD+ssNS(+) | | | | | 132 | | 139 | | | 152 | 156 | | | | 163 | 165 | | 167 | | | | | | | | | | |
| >N-NTD+dsNS | | | | | | | | | | 152 | 156 | | | | 163 | 165 | 166 | 167 | 170 | 172 | 173 | | 175 | | | | | |
| consensus | | | | | | | | | | * | * | | | | : | : | : | : | | : | | | | | | | | |
| >N-NTD-SR+TRS(-) | | | | | | | | | | 152 | | 157 | | | | 165 | 166 | | | 172 | 173 | 174 | | | | | | |
| >N-NTD-SR+TRS(+) | | | | | | | | | | 152 | | | | | 163 | 165 | 166 | | | | 173 | 174 | | 176 | | | | |
| >N-NTD-SR+dsTRS | | | | | | | | | | 152 | | | | | | | 166 | | | | | | | | 177 | | 197 | |
| consensus | | | | | | | | | | * | | | | | | : | * | | | | : | : | | | | | | |
| >N-NTD-SR+ssNS(-) | 112 | | 118 | 131 | | | | | | 152 | | | | | | | 166 | | | 172 | 173 | | | 176 | | | | 210 |
| >N-NTD-SR+ssNS(+) | 112 | | 118 | 131 | | | | | | 152 | | | | | | | 166 | | | 172 | 173 | | | 176 | | | | 210 |
| >N-NTD-SR+dsNS | # | # | | 131 | | | | # | # | 152 | | # | 159 | | 163 | | 166 | | # | # | 173 | # | # | 176 | | 196 | 197 | |
| consensus | * | | : | * | | | | | | * | | | | | | | * | | | * | * | | | * | | | | : |

**Figure S6.** Alignment of the consensus residues with CSP values higher than $\Delta\delta_{ave}$ + SD for the interaction of N-NTD and N-NTD-SR with ssDNAs and dsDNAs identified in the NMR titration experiments. The asterisk (*) and colon (:) indicate fully (three) and partial (two) consensus residues, respectively. Residue with resonance signal broadened beyond detection upon DNA binding is represented by the hashtag (#). The symbol § denotes the residues of N-NTD reported by Dinesh and cols. (2020) for the interaction with 7-nucleotide and 10-nucleotide ssTRS(+) RNA (5'–CUAAACG–3', 5'–UCUCUAAACG–3'), and 7-nucleotide non-specific RNA duplex (5'–CACUGAC–3' and 5'–GUCAGUG–3') (1). Numbers represent the amino acid position in the protein sequence.

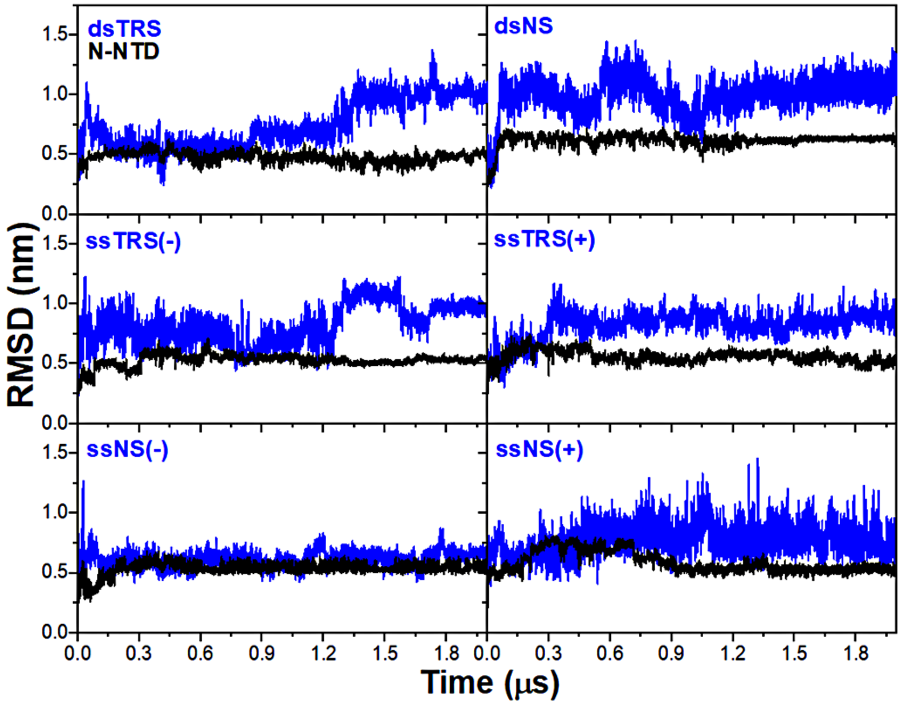


**Figure S7.** RMSD values of the backbone atoms of N-NTD (black) and protein-bound DNA (blue) along the 2 μs MD simulations.

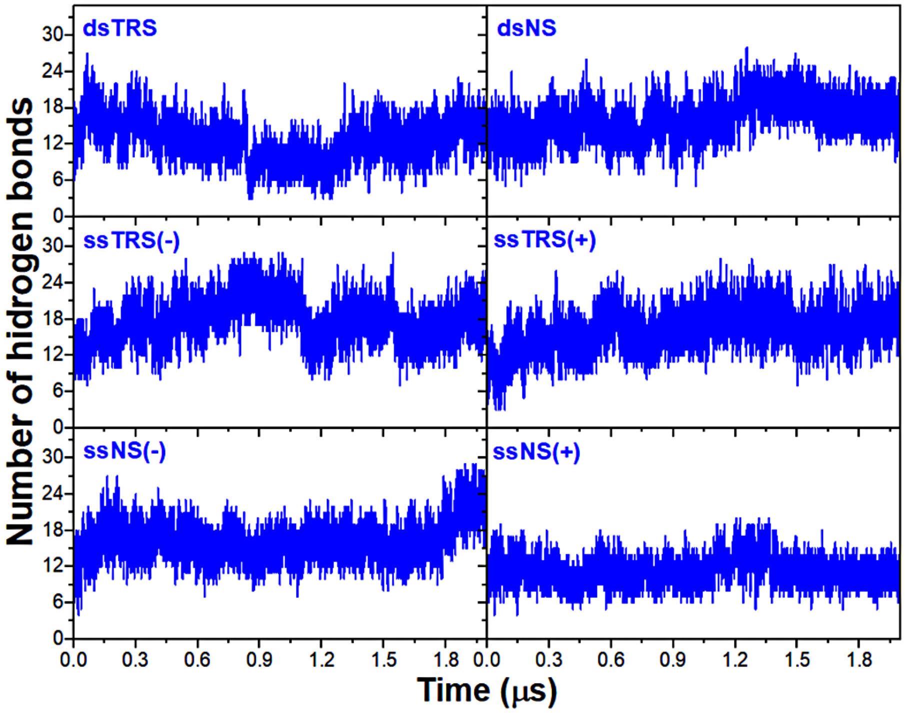


**Figure S8.** Number of protein-DNA hydrogen bonds formed between N-NTD and DNAs along the 2 μs MD simulations.

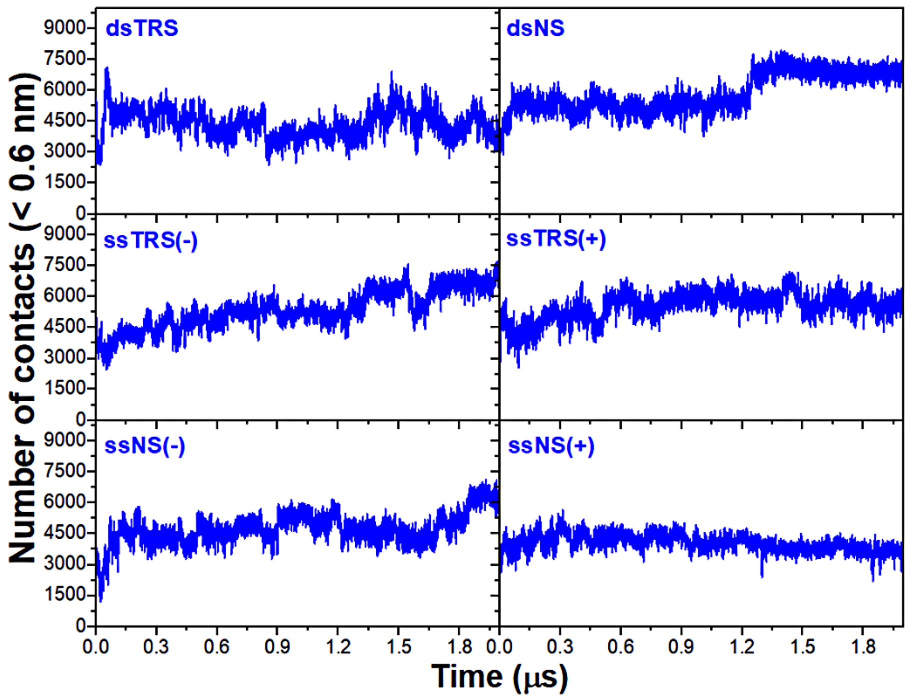


**Figure S9.** Number of contacts < 0.6 nm between the N-NTD and DNA atoms for the along the 2 μs MD simulations.

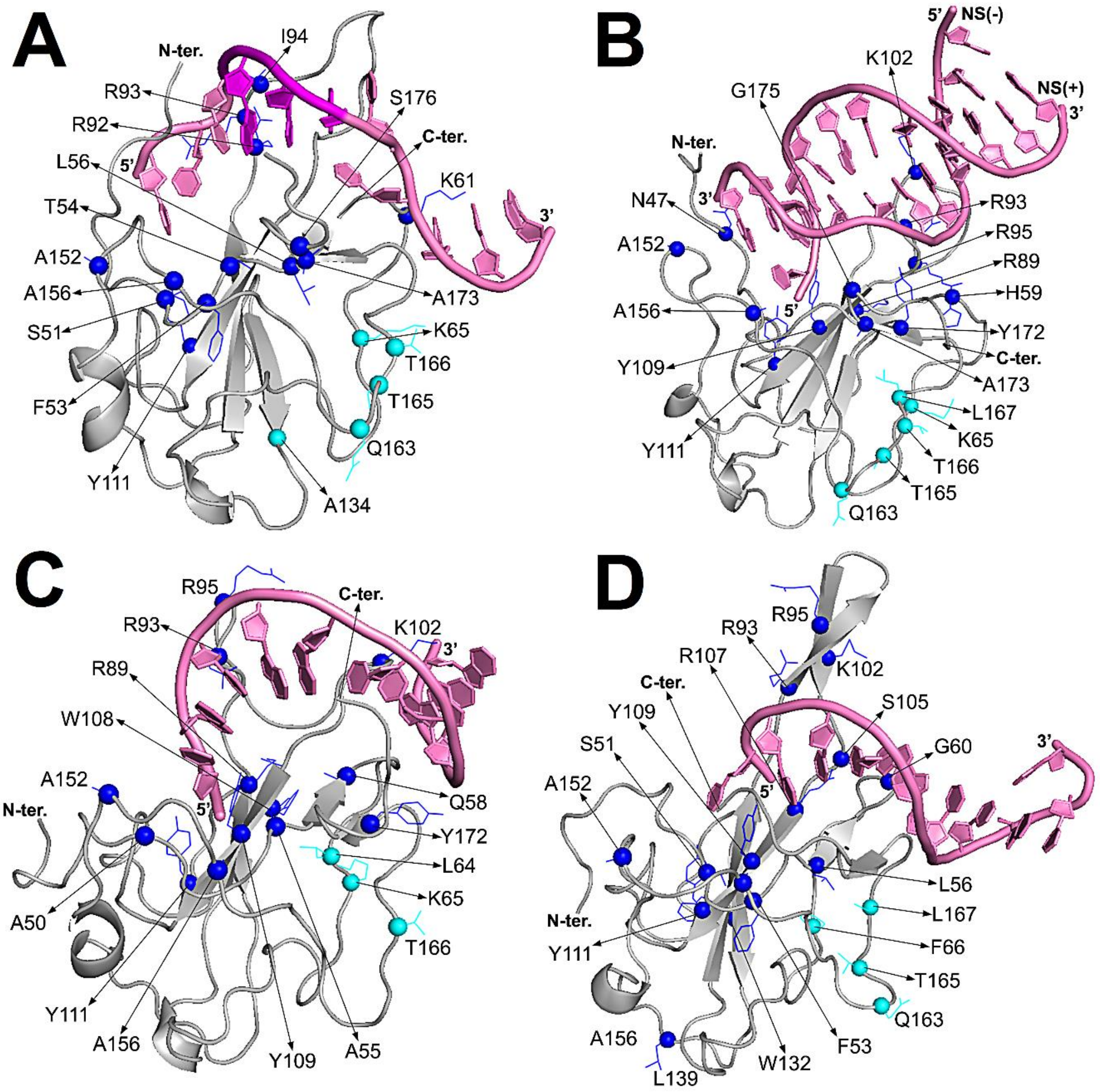


**Figure S10.** Representative structure models of the N-NTD complexed with (A) ssTRS(+), (B) dsNS, (C) ssNS(–), and (D) ssNS(+) obtained from the molecular docking and molecular dynamic simulations. The spheres denote the residues with CSP values higher than $\Delta\delta_{ave}$ + SD, participating directly in the binding interface (blue) and in the remote region (cyan). The proteins are shown as a cartoon model, DNAs are represented as a cartoon-ring model, and the TTT motif in TRS is colored in magenta.

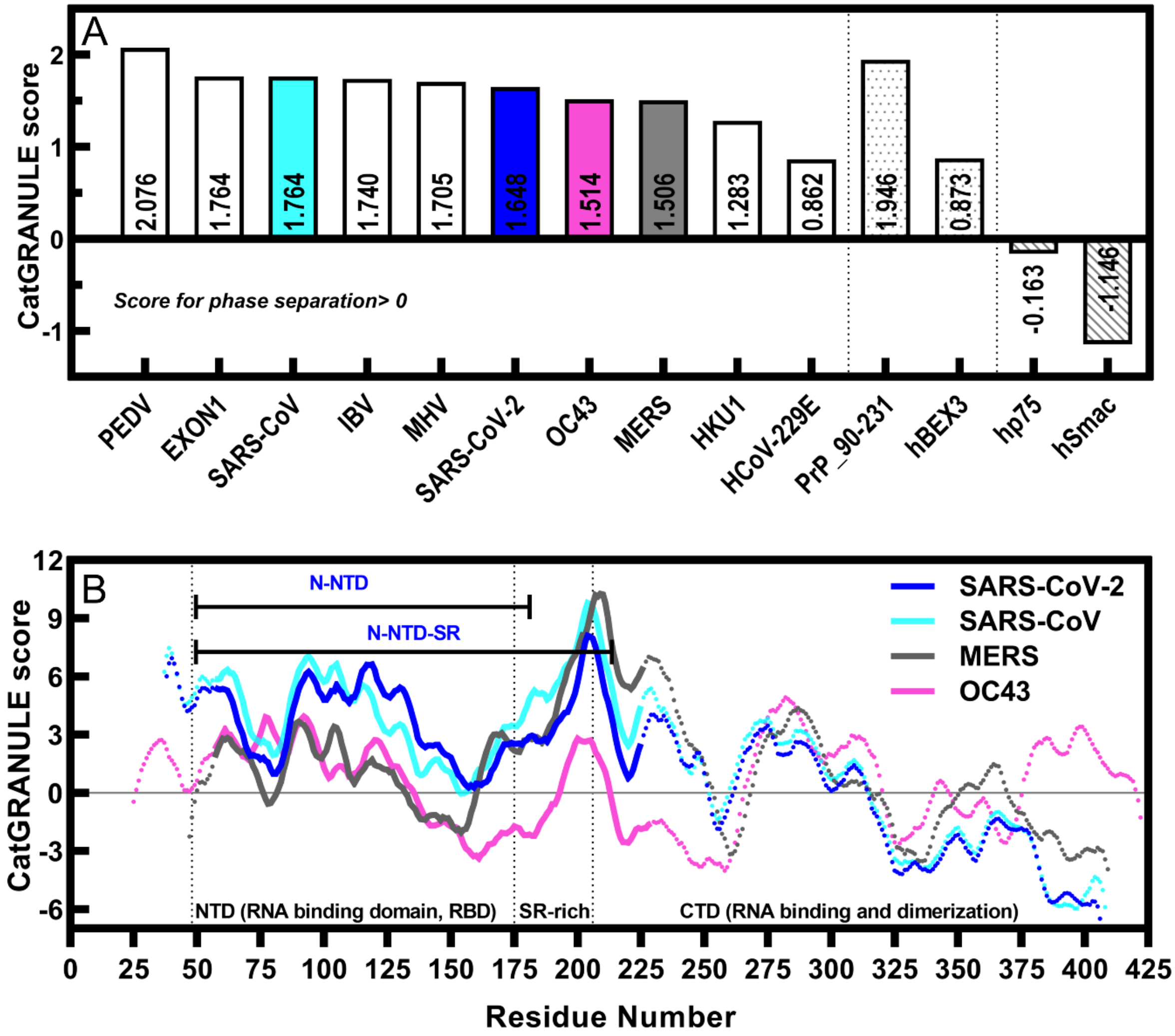


**Figure S11. The liquid-liquid phase separation of N-protein is predicted to be conversed between coronaviruses and the SR-rich region shows the highest propensity. (A)** Scores for liquid-liquid phase separation, based on catGRANULE algorithm (2) (score > 0 indicates granule formation; above 1.0 denotes strong propensity; available at http://service.tartaglialab.com/new_submission/catGRANULE). Scores of N-protein from SARS-Cov-2 (Uniprot ID P0DTC9); SARS-CoV (P59595); MERS (R9UM87); OC43 (P33469); HCoV-229E (P15130); PEDV (Q07499); IBV (P69596); HKU1 (Q5MQC6); EXON1 (D2E1E2) were compared with controls. Positive controls composed by proteins that were experimentally proved to undergo *in vitro* liquid-liquid phase separation modulated by nucleic acids, such as the prion protein (PrP_90-231; Uniprot ID P04925) (3) and Brain Expressed-X-linked 3 (BEX3; Uniprot ID Q00994) (4) represented by dotted bars. Negative controls composed by folded proteins such as hp75 (Uniprot ID P02768) and hSmac/Diablo (Uniprot ID Q9NR28) represented by stripped bars. Betacoronaviruses (colored bars) had phase separation propensity calculated along primary structure in **(B)** by catGRANULE. The dotted lines mark the domain organization of N-protein (1-48 is the IDR; 48-175 is the RBD; 176-206 is the SR-rich region followed by the C-terminus). The N-terminal constructs studied herein are highlighted in blue font. The SR-rich region, which is only present in the

N-NTD-SR construct, is likely to majorly contribute to N-protein liquid-liquid phase separation.

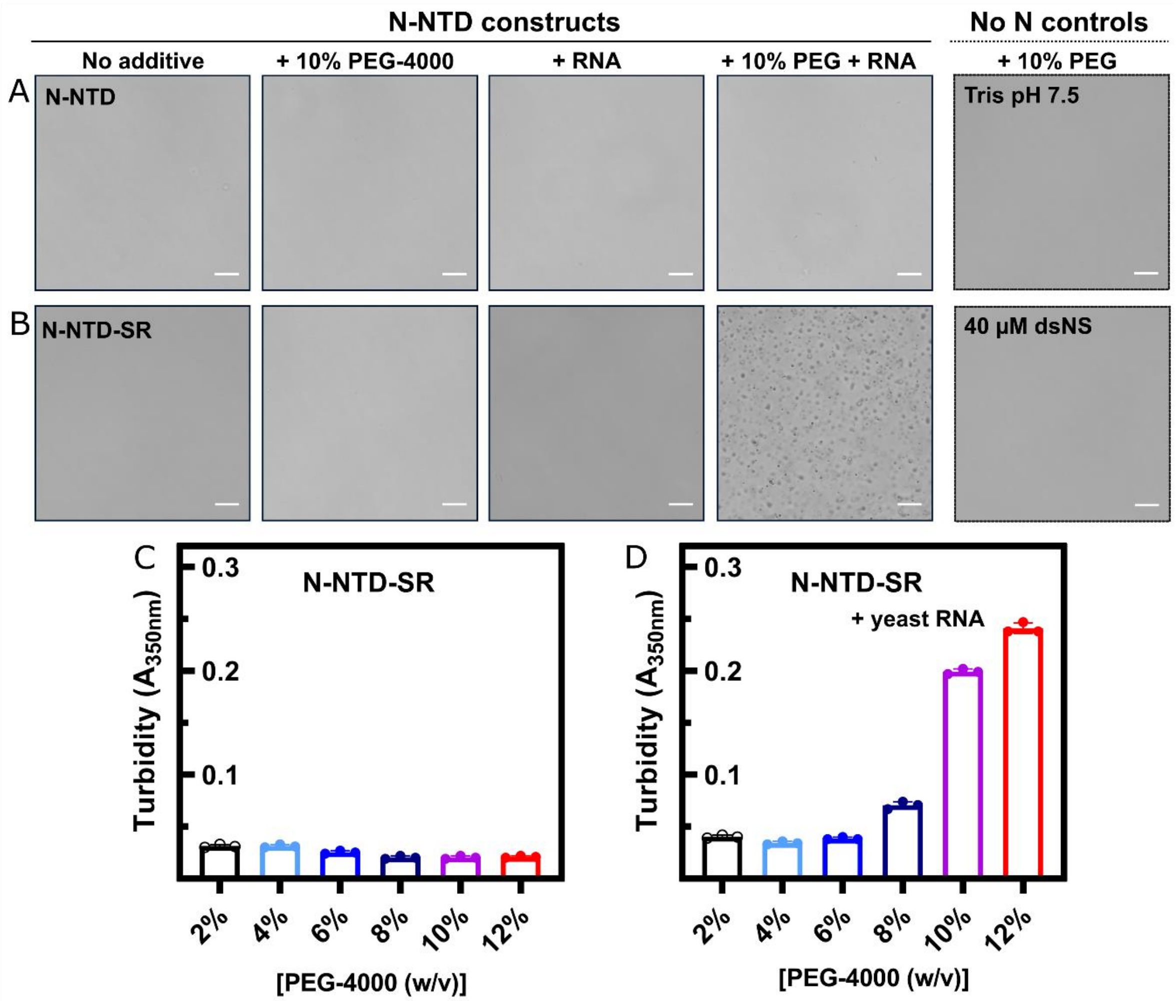


**Figure S12. The N-protein SR-rich region is important for N-NTD demixing with RNA.** Representative phase contrast micrographs of 20 µM N-NTD (A) and 20 µM N-NTD-SR (B) in 20 mM Tris-HCl buffer (pH 7.5) containing 30 mM NaCl. Left to right: only protein in buffer; protein in buffer containing the crowding agent 10% (w/v) PEG-4000; protein in buffer + Torula yeast RNA (1:2); protein in buffer + 10% (w/v) PEG-4000 + Torula yeast RNA (1:2). Controls are shown in dashed frames. Only vehicle consisting of buffer + 10% (w/v) PEG-4000. Only nucleic acid in the highest concentration used in the experiments (40 µM ds22-bp). Phase separation of 20 µM N-NTD-SR in 20 mM Tris-HCl buffer (pH 7.5) containing 30 mM NaCl monitored by absorbance measurements at 350 nm as a function of increasing concentrations of PEG-4000 (2% (black), 4% (light blue), 6% (marine blue), 8% (dark blue), 10% (purple), 12% (red)) in the absence (C) and presence (D) of Torula yeast RNA extract.

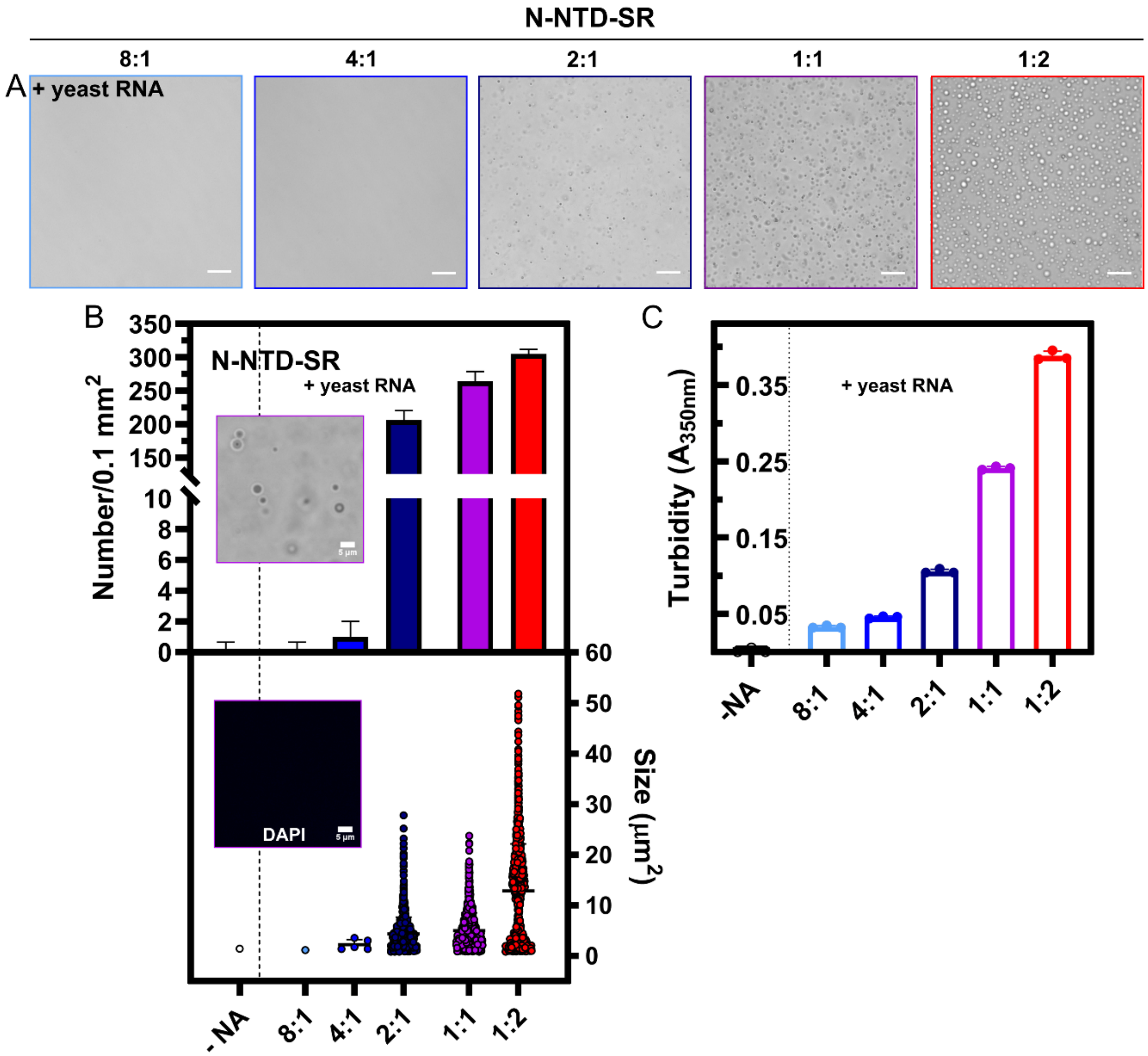


**Figure S13. Phase separation is promoted by increasing concentrations of yeast RNA.** LLPS assessment of 20 µM N-NTD-SR in the presence of increasing Torula yeast RNA concentrations: 8:1 (light blue); 4:1 (marine blue); 2:1 (dark blue); 1:1 (purple); 1:2 (red) protein:RNA molar ratios, respectively, in 20 mM Tris-HCl buffer (pH 7.5) containing 30 mM NaCl. (A) Representative phase contrast micrographs. (B) Top graph: Mean number of condensates ± S.D. per 100 $\mu m^2$ area (n= 5 images). Top inset: Phase contrast microscopy in the presence of DAPI for 1:1 protein:RNA stoichiometry. Bottom graph: Scatter plot from the size of condensates (represented as area in $\mu m^2$) obtained from micrographs analysis. The following number of condensates had their size measured: 8:1 (N= 1); 4:1 (N= 5); 2:1 (N= 1028); 1:1 (N= 1321) and 1:2 (N= 1753). Bottom inset: corresponding DAPI emission (0.25 µg/mL) from the top graph inset image (no fluorescence observed). All conditions contained 10% (w/v) PEG-4000. Scale bar, 20 µm apart from insets (5 µm). (C) Phase separation of 20 µM N-NTD-SR in 20 mM Tris-HCl buffer (pH 7.5) containing 30 mM NaCl) in the presence of 10% (w/v) PEG-4000 monitored by absorbance measurements at 350 nm as a function of increasing concentrations of Torula yeast RNA extract. The protein:RNA stoichiometries are the same as in (A).

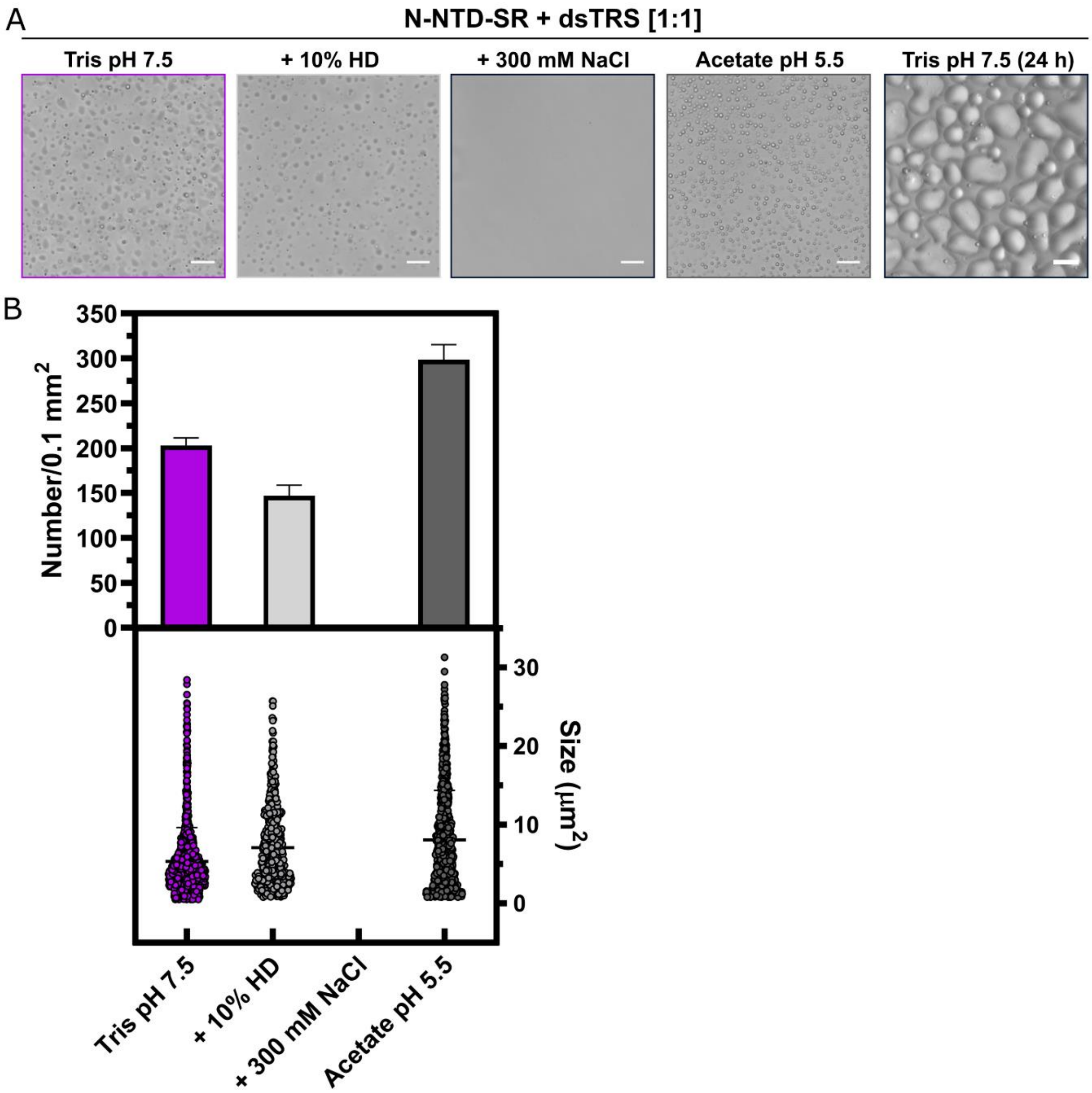


**Figure S14. N-NTD-SR:dsTRS liquid-liquid phase separation is driven by electrostatic contacts and promoted by acidic pH.** (A) Representative phase contrast micrographs from samples containing 20 µM N-NTD-SR in the presence of 20 µM dsTRS (1:1 molar ratio). Top left: sample in 20 mM Tris-HCl buffer (pH 7.5) containing 30 mM NaCl). Top right: with addition of 10% (w/v) 1,6-hexanediol (HD). Bottom left: with addition of 300 mM NaCl. Bottom right: sample in 20 mM sodium acetate buffer pH 5.5 with 30 mM NaCl. (B) Quantification of N-NTD-SR:dsTRS condensates in the presence of additives. Top graph: Mean number of condensates ± S.D. per 100 $\mu m^2$ area (n= 5 images). Bottom graph: Scatter plot from condensates' size (represented as area in $\mu m^2$) obtained from micrographs analysis. The following number of condensates' size were analyzed: sample in buffer (purple data, N= 1013); sample in buffer containing 10% HD (light gray data, N=598); no condensates visualized in presence of 300 mM NaCl (absence of column, N=0); sample in 20 mM sodium acetate pH 5.5, 30 mM NaCl (dark gray column, N=1174). All conditions contained 10% (w/v) PEG-4000. Scale bar, 20 µm.

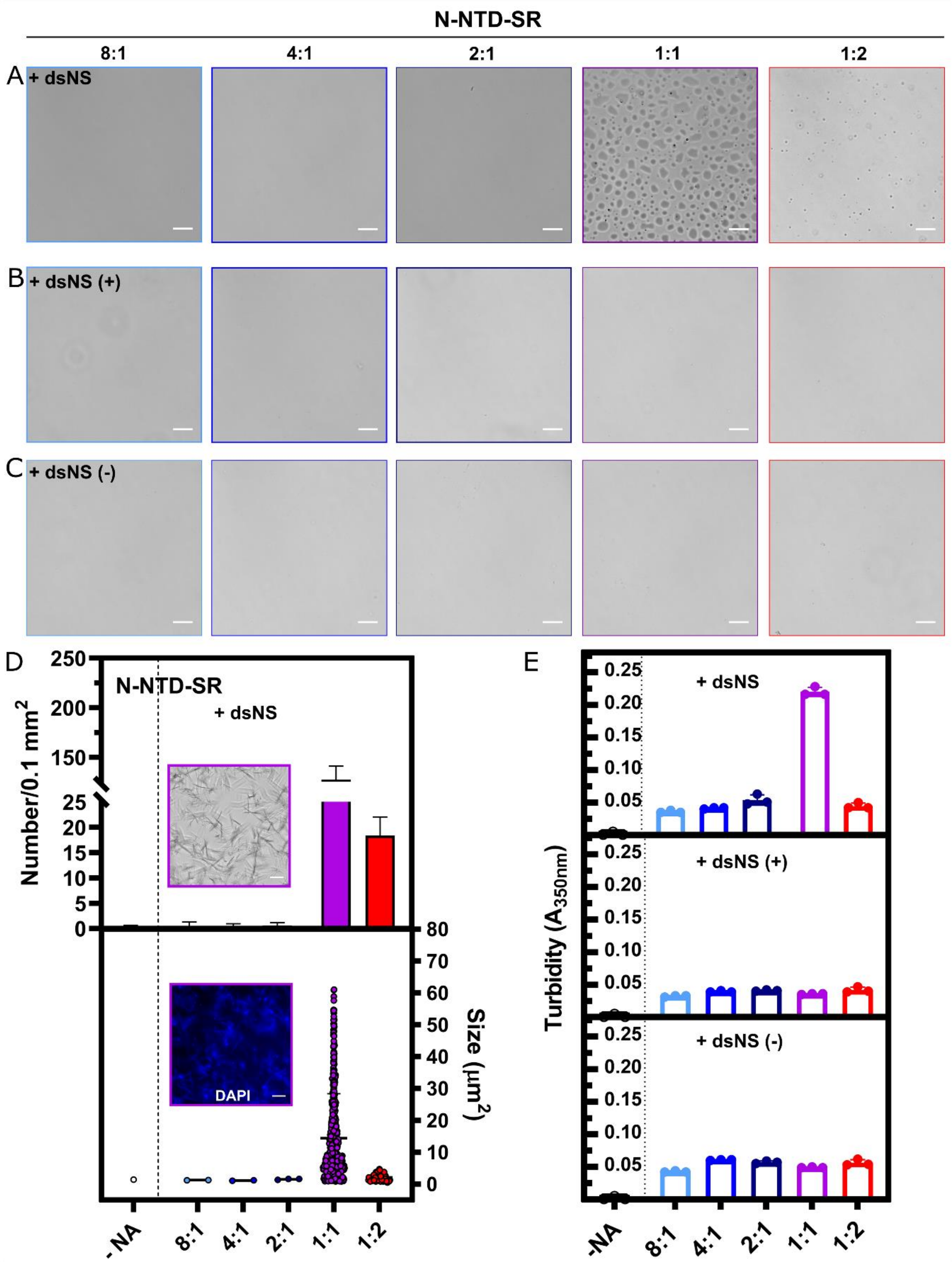


**Figure S15. The non-specific DNA oligonucleotide needs secondary structure to induce N-NTD-SR liquid-liquid phase separation.** Representative phase contrast micrographs of 20 µM N-NTD-SR in the presence of dsNS (A); ssNS(+) (B) or ssNS(–) (C) at the following protein:DNA stoichiometries: 8:1 (light blue); 4:1 (marine blue); 2:1 (dark blue); 1:1

(purple); 1:2 (red), respectively, in 20 mM Tris-HCl buffer (pH 7.5) containing 30 mM NaCl. (D) Top graph: Mean number of condensates ± S.D. per 100 μm$^2$ area (n= 5 images). Top inset: Phase contrast microscopy in the presence of DAPI for 1:1 N-NTD-SR:dsNS). Bottom graph: Scatter plot from the size of condensates (represented as area in μm$^2$) obtained from micrographs analysis. Since no phase separation was observed for the single-stranded oligonucleotides, the following number of condensates' size were measured for the dsNS: -NA (1); at 8:1 (N= 2); 4:1 (N= 2); 2:1 (N= 3); 1:1 (N= 632) and 1:2 (N= 81). Bottom inset: corresponding DAPI emission from the top graph inset image acquired after 2 h incubation evidencing needle-like crystals formed by N-NTD-SR:dsNS at 1:1 stoichiometry. All conditions contained 10% (w/v) PEG-4000. Scale bar, 20 μm including insets. (E) Phase separation of 20 μM N-NTD-SR in 20 mM Tris-HCl buffer (pH 7.5) containing 30 mM NaCl in the presence of 10% (w/v) PEG-4000 monitored by absorbance measurements at 350 nm as a function of increasing concentrations of the non-specific DNA oligonucleotides dsNS (top graph), ssNS(+) (middle graph), and ssNS(–) (bottom graph). The protein:DNA stoichiometries are the same as in (A).

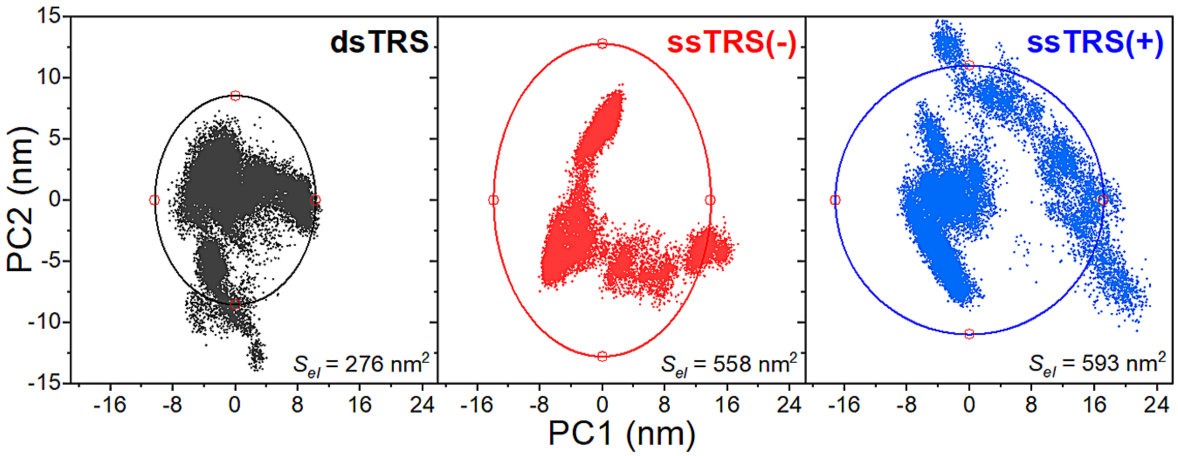


**Figure S16.** PCA scatter plots PC1 and PC2 for N-NTD complexed with dsTRS (black dots, *left*), ssTRS(–) (red dots, *middle*), and ssTRS(+) (blue dots, *right*). The extent of the conformational space for each scatter plot was measured by fitting an elliptical shell (solid lines) that contains 95% (confidence) of the density. The area of the ellipse ($S_{el}$) is a measure of the conformational space for the free and bound protein. The N-NTD complexed with dsTRS explored a smaller conformational space along the 2 μs MD simulation, meaning a more restricted motion of the protein, which may be a result of a better-defined position and orientation of the DNA duplex.

```
                                                               NLS1
OC43        MSFTPGKQSSSRASSGNRS-VNGILKWADQSDQFRNVQTRGRRAQPKQTATSQQPSGGNV
HKU1        MSYTPGHYAGSRSSSGNRSGILKKTSWADQSERNYQTFNRGRKTQPKFTVSTQPQ--GNT
SARS-CoV2   ---------MSDNGPQ-NQRNAPRITFGGPSDSTGSNQNGERSG----ARSKQRRPQGLP
SARS        ---------MSDNGPQSNQRSAPRITFGGPTDSTDNNQNGGRNG----ARPKQRRPQGLP
MERS        ---------------MASPAAPRAVSFADNNDITNTNLSRGRG--------RNPKPRAAP
                                     .:.. .:      .  *          :    .

OC43        VPYYSWFSGITQFQKGKEFEFAEGQGVPIAPGVPATEAKGYWYRHNRRSFKTADGNQRQL
HKU1        IPHYSWFSGITQFQKGRDFKFSDGQGVPIAFGVPPSEAKGYWYRHSRRSFKTADGQQKQL
SARS-CoV2   NNTASWFTALTQHGK-EDLKFPRGQGVPINTNSSPDDQIGYYRRATRR-IRGGDGKMKDL
SARS        NNTASWFTALTQHGK-EELRFPRGQGVPINTNSGPDDQIGYYRRATRR-VRGGDGKMKEL
MERS        NNTVSWYTGLTQHGK-VPLTFPPGQGVPLNANSTPAQNAGYWRRQDRK-INTGNG-IKQL
                **::.:**. *   : *. *****:  .  . :  **: *  *: .. .:*  ::*

OC43        LPRWYFYYLGTGPHAKDQYGTDIDGVYWVASNQADVNTPADIVDRDPSSDEAIPTRFPPG
HKU1        LPRWYFYYLGTGPYANASYGESLEGVFWVANHQADTSTPSDVSSRDPTTQEAIPTRFPPG
SARS-CoV2   SPRWYFYYLGTGPEAGLPYGANKDGIIWVATEGALNTPKDHIGTRNPANNAAIVLQLPQG
SARS        SPRWYFYYLGTGPEASLPYGANKEGIVWVATEGALNTPKDHIGTRNPNNNAATVLQLPQG
MERS        APRWYFYYTGTGPEAALPFRAVKDGIVWVHEDGATDAP-STFGTRNPNNDSAIVTQFAPG
             ******* **** *   :    :*: **  . *   .   .  *:* .: *   ::. *

OC43        TVLPQGYYIEGS-GRSAPNSRSTSRTSSRASSAGSRSRANSGNRTPTSGVTPDMADQIAS
HKU1        TILPQGYYVEGS-GRSASNSRPGSRSQSRGPNTRSLSRSNSNFRHSDSIVKPDMADEIAN
SARS-CoV2   TTLPKGFYAEGSRGGSQASSRSSSRSRNSSRNSTPGSSRGTSP---ARMAGNGGDAALAL
SARS        TTLPKGFYAEGSRGGSQASSRSSSRSRGNSRNSTPGSSRGNSP---ARMASGGGETALAL
MERS        TKLPKNFHIEGTGGNSQSSSRASSLSRNSSRSSSQGSRSGNSTRGTSPGPSGIGAVGGDL
            * **:.:: **: * * ..**. * : . . .:   *  ...
              NES                               NLS2
OC43        LVLAKLGKDATKPKQVTKHT-----AKEVRQKILNKPRQKRSPNKQCTVQQCFGKRGP--
HKU1        LVLAKLGKDS-KPQQVTKQN-----AKEIRHKILTKPRQKRTPNKHCNVQQCFGKRGP--
SARS-CoV2   LLLDRLNQLESKMSGKGQQQQGQTVTKKSAAEASKKPRQKRTATKAYNVTQAFGRRGPEQ
SARS        LLLDRLNQLESKVSGKGQQQQGQTVTKKSAAEASKKPRQKRTATKQYNVTQAFGRRGPEQ
MERS        LYLDLLNRLQALESGKVKQSQPKVITKKDAAAAKNKMRHKRTSTKSFNMVQAFGLRGPGD
            * *  *.:     .   ::      :*:      .* *:**:..*  .: *.** ***

OC43        -NQNFGGGEMLKLGTSDPQFPILAELAPTAGAFFFGSRLELAKVQNLSGNPDEPQKDVYE
HKU1        -SQNFGNAEMLKLGTNDPQFPILAELAPTPGAFFFGSKLELVKRE---SEADSPVKDVFE
SARS-CoV2   TQGNFGDQELIRQGTDYKHWPQIAQFAPSASAFFGMSRIGMEVTPSG-----------TW
SARS        TQGNFGDQDLIRQGTDYKHWPQIAQFAPSASAFFGMSRIGMEVTPSG-----------TW
MERS        LQGNFGDLQLNKLGTEDPRWPQIAELAPTASAFMGMSQFKLTHQNNDDHGNP-----VYF
             . ***. :: : **.  ::* :*::**:..**:  *:: :
                                                                NLS3
OC43        LRYNGAIRFDSTLSGFETIMKVLSENLNAYQ---QQDGMMNMSPKPQRQRGHKNGQGEND
HKU1        LRYSGSIRFDSTLPGFETIMKVLKENLDAYVNSNQNTVSGSLSPKPQRKRGVKQSPELFD
SARS-CoV2   LTYTGAIKLDDKDPNFKDQVILLNKHIDAYK-----------TFPPTEPKK--DKKKKAD
SARS        LTYHGAIKLDDKDPQFKDNVILLNKHIDAYK-----------TFPPTEPKK--DKKKKTD
MERS        LRYSGAIKLDPKNPNYNKWLELLEQNIDAYK-----------TFPKKEKKQKAPKEESTD
            * * *:*::* . . ::  : :*.::::**            :    . :         *
                NLS3
OC43        NISVAVPKSRVQQNKSIELTAEDISLLKKMDEPYTEDTSEI---
HKU1        SLNLSADTQHISN----DFTPEDHSLLATLDDPYVEDSVA----
SARS-CoV2   ETQALPQRQKKQQTVTLLPAADLDDFSKQLQQSMSSADSTQA--
SARS        EAQPLPQRQKKQPTVTLLPAADMDDFSRQLQNSMSGASADSTQA
MERS        QMSEPPKEQRVQGSITQRTRTRPSVQPGPMIDVNTD--------
            . .     .: .        .        : :
```

**Figure S17.** Sequence alignment of the N protein from SARS-CoV-2, SARS-CoV, CoV-MERS, CoV-HKU1, and CoV-OC43 performed using the webserver ClustalW (5). Sequence homology between hCoV N protein is denoted by asterisk (*), colon (:), and period (.). The asterisk indicates fully conserved residue, colon represents conservation between groups of strongly similar properties, and period indicates conservation between groups of weakly similar properties. The residue sequences of the SARS-CoV-2 N-NTD and its SR-motif are denoted in red and blue, respectively. The nuclear localization signals are indicated by NLS1, NLS2, and NLS3 with the residue sequence for SARS-CoV-2 and SARS-CoV highlighted in cyan and gray, respectively. The nuclear export signal is indicated by NES with the residue sequence for SARS-CoV-2 and SARS-CoV highlighted in magenta and gray, respectively. The residue E174 is denoted in green.

**Table S1.** Persistent hydrogen bonds (and salt bridges) between N-NTD and dsTRS.

| dsTRS | | | | |
|---|---|---|---|---|
| **Donor** | **Donor Atom** | **Acceptor** | **Acceptor atom** | **Persistence percentage** |
| T1(+) | O5' | ASN48 | OD1 | 14.579 |
| ASN48 | ND2 | T1(+) | O4' | 14.199 |
| ARG88 | NH1 | C2(+) | O2P | 56.627 |
| ASN48 | ND2 | C2(+) | O1P | 42.968 |
| ARG88 | NH2 | C2(+) | O2P | 20.094 |
| ARG88 | NH2 | C2(+) | O1P | 18.484 |
| C2(+) | N4 | GLU174 | OE2 | 15.709 |
| C2(+) | N4 | GLU174 | OE1 | 12.979 |
| ASN47 | N | C2(+) | O1P | 10.274 |
| ARG107 | NH1 | T3(+) | O4 | 49.028 |
| ARG92 | NE | T3(+) | O2P | 12.009 |
| ARG107 | NH2 | T3(+) | O4 | 10.324 |
| ARG92 | NH2 | A4(+) | O2P | 10.174 |
| ARG95 | NH1 | A5(+) | O2P | 11.854 |
| GLY97 | N | G4(-) | O2P | 31.763 |
| LYS102 | NZ | G4(-) | O2P | 26.994 |
| LYS102 | NZ | G4(-) | O1P | 18.974 |
| LYS102 | NZ | G4(-) | N7 | 12.384 |
| GLY97 | N | G4(-) | O1P | 11.399 |
| LYS61 | NZ | G5(-) | O2P | 40.308 |
| LYS102 | NZ | G5(-) | O6 | 11.439 |
| GLY60 | N | T6(-) | O2P | 32.713 |
| GLY60 | N | T6(-) | O1P | 30.848 |
| SER105 | OG | T6(-) | O2P | 22.159 |
| HIS59 | NE2 | T7(-) | O1P | 38.308 |
| ARG177 | NH2 | T7(-) | O2P | 22.799 |
| TYR172 | OH | T7(-) | O2P | 21.344 |
| ARG177 | NH1 | T7(-) | O1P | 11.399 |
| ARG177 | NE | T7(-) | O1P | 10.509 |
| ARG107 | NH2 | T8(-) | O4 | 36.593 |
| ARG177 | NH2 | T8(-) | O2P | 16.424 |
| ARG177 | NH1 | T8(-) | O2P | 13.154 |
| GLY178 | N | T8(-) | O2P | 11.609 |
| ARG177 | NE | T8(-) | O1P | 11.309 |
| A9(-) | N6 | GLU174 | OE2 | 14.734 |
| G9(-) | N2 | ALA42 | O | 16.694 |

**Table S2.** Persistent hydrogen bonds (and salt bridges) between N-NTD and dsNS.

| dsNS | | | | |
|---|---|---|---|---|
| **Donor** | **Donor Atom** | **Acceptor** | **Acceptor atom** | **Persistence percentage** |
| TYR111 | OH | C1(+) | O5' | 87.506 |
| ASN48 | ND2 | C1(+) | O2 | 74.066 |
| ASN48 | ND2 | A2(+) | O4' | 79.506 |
| A2(+) | N6 | TYR109 | OH | 46.253 |
| ARG88 | NH2 | A2(+) | O1P | 44.448 |
| ARG88 | NH1 | A2(+) | O1P | 40.648 |
| ARG88 | NH1 | A2(+) | O2P | 34.933 |
| LEU45 | N | A2(+) | N3 | 27.964 |
| ASN47 | ND2 | A2(+) | N3 | 17.499 |
| ARG92 | NH2 | C3(+) | O2P | 19.569 |
| ARG92 | NH1 | C3(+) | O2P | 16.679 |
| ARG92 | NE | C3(+) | O2P | 10.604 |
| ALA42 | N | T4(+) | O2 | 19.659 |
| LYS102 | N | G4(-) | O2P | 55.752 |
| LYS102 | NZ | G4(-) | N7 | 17.869 |
| ARG93 | NH2 | G5(-) | O2P | 80.041 |
| ARG93 | NH1 | G5(-) | O2P | 67.972 |
| LYS102 | NZ | G5(-) | O6 | 48.863 |
| MET101 | N | G5(-) | O2P | 44.918 |
| LYS100 | NZ | G5(-) | O1P | 22.094 |
| ARG93 | NH2 | G5(-) | O5' | 21.444 |
| GLY99 | N | G5(-) | O1P | 17.599 |
| SER105 | OG | T6(-) | O2P | 47.303 |
| SER105 | N | T6(-) | O2P | 42.653 |
| ARG177 | NH2 | T6(-) | O1P | 27.444 |
| ARG93 | NH1 | G5(-) | O1P | 19.864 |
| ARG177 | NH1 | T6(-) | O1P | 17.459 |
| LYS102 | NZ | T6(-) | O4 | 11.899 |
| ARG177 | NH1 | C7(-) | O2P | 34.583 |
| SER105 | OG | C7(-) | N4 | 22.519 |
| ARG177 | NH2 | C7(-) | O2P | 15.014 |
| TYR172 | OH | C7(-) | O2P | 10.309 |
| ARG107 | NH1 | G9(-) | O6 | 55.462 |
| G9(-) | N2 | ALA42 | O | 34.598 |
| ARG107 | NH2 | G9(-) | O6 | 17.359 |
| GLY44 | N | T10(-) | O2 | 32.723 |
| G11(-) | N2 | LEU45 | O | 36.778 |
| G11(-) | O3' | GLY44 | O | 24.269 |
| G11(-) | N2 | ASN47 | OD1 | 21.884 |

**Table S3.** Persistent hydrogen bonds (and salt bridges) between N-NTD and ssTRS(–).

| ssTRS(-) | | | | |
|---|---|---|---|---|
| **Donor** | **Donor Atom** | **Acceptor** | **Acceptor atom** | **Persistence percentage** |
| ASN47 | ND2 | C1 | O2 | 39.168 |
| C1 | N4 | SER180 | OC2 | 37.868 |
| C1 | N4 | SER180 | OC1 | 32.798 |
| C1 | O5' | THR49 | O | 20.004 |
| THR49 | OG1 | C1 | O5' | 18.069 |
| TYR111 | OH | C1 | O5' | 14.929 |
| TYR109 | OH | G2 | O2P | 81.581 |
| ALA42 | N | G2 | O1P | 34.533 |
| G2 | N1 | SER180 | OC2 | 24.599 |
| G2 | N1 | SER180 | OC1 | 22.829 |
| SER180 | OG | G2 | N1 | 20.089 |
| G2 | N2 | ARG177 | O | 19.794 |
| LEU45 | N | G2 | O1P | 18.994 |
| G2 | N2 | SER180 | OC1 | 18.649 |
| G2 | N2 | SER180 | OC2 | 15.734 |
| G2 | N2 | GLY178 | O | 12.084 |
| ARG149 | NH2 | G2 | O6 | 11.744 |
| ARG92 | NH2 | G2 | O2P | 11.049 |
| ARG107 | NH2 | C3 | O2P | 48.448 |
| ARG177 | N | C3 | O4' | 31.698 |
| ARG107 | NH1 | C3 | O2P | 30.493 |
| ARG93 | NH2 | C3 | O1P | 25.624 |
| ARG177 | NE | C3 | O2 | 17.514 |
| ARG177 | NH2 | C3 | O2P | 14.899 |
| ARG93 | NE | C3 | O1P | 14.064 |
| ARG177 | NH1 | C3 | O5' | 13.159 |
| ARG95 | NH1 | G4 | O2P | 35.548 |
| ARG95 | NH2 | G4 | O2P | 33.258 |
| ARG93 | NH2 | G4 | O1P | 13.449 |
| ARG107 | NH2 | G4 | N7 | 11.914 |
| ARG93 | NH2 | G4 | O2P | 11.859 |
| ARG93 | NE | G4 | O1P | 11.014 |
| ARG92 | NH2 | G5 | O6 | 31.083 |
| ARG95 | NE | G5 | O2P | 30.638 |
| G5 | N1 | GLU174 | OE1 | 23.599 |
| G5 | N1 | GLU174 | OE2 | 23.094 |
| ARG92 | NH1 | G5 | O6 | 23.054 |
| G5 | N2 | GLU174 | OE1 | 20.434 |
| ARG95 | NH2 | G5 | O5' | 20.164 |
| G5 | N2 | GLU174 | OE2 | 19.564 |
| ARG95 | NH2 | G5 | O2P | 14.729 |
| SER105 | N | T6 | O4 | 29.894 |
| SER105 | OG | T6 | N3 | 17.864 |
| TYR172 | OH | T6 | O2 | 10.819 |
| TYR172 | OH | T7 | O2 | 34.293 |
| GLY170 | N | A9 | O1P | 49.623 |

| | | | | |
|---|---|---|---|---|
| LYS169 | NZ | G10 | O1P | 47.848 |
| LYS61 | N | A11 | N1 | 14.344 |

**Table S4.** Persistent hydrogen bonds (and salt bridges) between N-NTD and ssTRS(+).

| **ssTRS(+)** | | | | |
|---|---|---|---|---|
| **Donor** | **Donor Atom** | **Acceptor** | **Acceptor atom** | **Persistence percentage** |
| GLY44 | N | T1 | O3' | 24.984 |
| T1 | N3 | ASN153 | OD1 | 11.039 |
| ARG107 | NH1 | C2 | O2 | 55.107 |
| GLY44 | N | C2 | O1P | 49.063 |
| ARG107 | NH2 | C2 | O2 | 46.193 |
| ARG93 | N | C2 | O3' | 30.173 |
| THR91 | N | C2 | N3 | 17.054 |
| ARG93 | NH1 | C2 | O2P | 15.149 |
| ARG93 | NH2 | C2 | O2P | 11.189 |
| ARG93 | NH2 | C2 | O1P | 10.979 |
| ARG92 | NE | T3 | O2P | 85.696 |
| ILE94 | N | T3 | O2P | 80.171 |
| ARG93 | N | T3 | O2P | 59.827 |
| ARG92 | NH2 | T3 | O2P | 53.822 |
| ARG93 | NE | T3 | O1P | 39.638 |
| ARG93 | NH2 | T3 | O1P | 29.774 |
| ARG92 | NH2 | T3 | O5' | 25.129 |
| MET43 | N | T3 | O2 | 13.219 |
| GLY41 | N | T3 | O3' | 11.714 |
| ARG92 | NH1 | A4 | N7 | 20.259 |
| GLY41 | N | A4 | O1P | 16.654 |
| A4 | N6 | GLU174 | OE2 | 15.519 |
| A4 | N6 | GLU174 | OE1 | 13.904 |
| ARG92 | NH2 | A4 | O2P | 10.359 |
| ARG177 | NH2 | A6 | O3' | 18.094 |
| ARG177 | NH1 | C7 | O1P | 26.609 |
| ARG177 | NH2 | C7 | O1P | 20.904 |
| TYR172 | OH | C7 | O2 | 10.769 |
| HIS59 | NE2 | C8 | O2P | 21.889 |
| LYS61 | N | G9 | O6 | 42.953 |
| GLY170 | N | G9 | O4' | 33.788 |
| HIS59 | NE2 | G9 | O5' | 29.959 |
| G9 | N1 | GLU62 | OE2 | 13.409 |
| G9 | N2 | GLU62 | OE2 | 12.664 |
| G9 | N2 | GLU62 | OE1 | 12.489 |
| G9 | N1 | GLU62 | OE1 | 12.439 |
| G9 | N2 | GLY60 | O | 10.804 |
| LYS169 | NZ | C10 | O1P | 36.403 |
| GLY170 | N | C10 | O1P | 23.959 |
| LYS169 | NZ | G11 | O1P | 16.139 |

**Table S5.** Persistent hydrogen bonds (and salt bridges) between N-NTD and ssNS(–).

| ssNS(-) | | | | |
|---|---|---|---|---|
| **Donor** | **Donor Atom** | **Acceptor** | **Acceptor atom** | **Persistence percentage** |
| C1 | O5' | ASN154 | O | 24.759 |
| ASN154 | ND2 | C1 | O2 | 12.644 |
| C1 | O5' | PHE53 | O | 11.774 |
| TYR109 | OH | G2 | O2P | 45.728 |
| THR91 | OG1 | G2 | O1P | 18.169 |
| THR91 | N | G2 | O1P | 14.094 |
| ARG88 | NH1 | G2 | O1P | 12.874 |
| ARG93 | NH2 | C3 | O2P | 30.998 |
| ARG95 | NH2 | C3 | O1P | 16.854 |
| ARG93 | NH1 | C3 | O2P | 13.529 |
| ARG95 | NH1 | C3 | O1P | 12.884 |
| ARG93 | NE | C3 | O2P | 11.774 |
| ARG107 | NH1 | G4 | O6 | 74.326 |
| ARG107 | NH2 | G4 | O6 | 68.737 |
| G4 | N2 | SER180 | OC1 | 42.148 |
| ARG93 | NH2 | G4 | O2P | 25.424 |
| G4 | N1 | SER180 | OC1 | 20.654 |
| ARG93 | NH1 | G4 | O2P | 19.449 |
| ARG93 | NH1 | G4 | N7 | 15.744 |
| G4 | N2 | SER180 | OC2 | 13.974 |
| ARG93 | NE | G4 | O2P | 13.334 |
| G4 | N1 | SER180 | OC2 | 12.244 |
| G5 | N2 | SER180 | OC2 | 64.952 |
| G5 | N2 | SER180 | OC1 | 30.078 |
| SER180 | N | T6 | O2 | 80.641 |
| SER180 | OG | T6 | O4' | 80.456 |
| ARG177 | NH1 | T6 | O2 | 11.934 |
| HIS59 | NE2 | C7 | O2 | 83.621 |
| ARG177 | NH1 | C7 | O2 | 68.262 |
| SER105 | OG | C7 | N4 | 43.693 |
| C7 | N4 | ASP103 | O | 17.694 |
| A8 | N6 | ASP103 | O | 16.174 |
| TYR172 | OH | G9 | O1P | 86.146 |
| G9 | N2 | ASP103 | OD2 | 36.903 |
| G9 | N2 | ASP103 | OD1 | 32.158 |
| LYS61 | N | T10 | O1P | 50.137 |
| LYS102 | NZ | T10 | O2 | 17.034 |
| LYS61 | NZ | G11 | O1P | 37.243 |
| LYS61 | NZ | G11 | O2P | 30.938 |
| ASP103 | N | G11 | O3' | 12.754 |

**Table S6.** Persistent hydrogen bonds (and salt bridges) between N-NTD and ssNS(+).

| ssNS(+) | | | | |
|---|---|---|---|---|
| **Donor** | **Donor Atom** | **Acceptor** | **Acceptor atom** | **Persistence percentage** |
| TYR111 | OH | C1 | N4 | 55.397 |
| C1 | N4 | ASN47 | O | 48.013 |
| TYR109 | OH | C1 | O5' | 38.443 |
| C1 | N4 | THR49 | O | 25.709 |
| SER176 | OG | A2 | N1 | 93.46 |
| A2 | N6 | SER176 | O | 92.245 |
| ARG92 | NE | A2 | O2P | 24.794 |
| ARG92 | NE | A2 | O1P | 20.869 |
| ARG92 | NH1 | A2 | O2P | 19.324 |
| ARG92 | NH2 | A2 | O2P | 17.249 |
| ARG92 | NH2 | A2 | O1P | 15.019 |
| SER176 | OG | A2 | N6 | 13.449 |
| ARG92 | NH1 | A2 | O1P | 11.304 |
| C3 | N4 | GLU174 | OE2 | 48.983 |
| C3 | N4 | GLU174 | OE1 | 45.353 |
| ARG92 | NH1 | C3 | O2P | 17.549 |
| ARG92 | NH2 | C3 | O2P | 15.644 |
| ARG107 | NH2 | T4 | O4 | 43.378 |
| TYR172 | OH | G5 | N2 | 68.177 |
| TYR172 | OH | A6 | N3 | 50.672 |
| GLY170 | N | C8 | O1P | 77.686 |
| LYS169 | NZ | G9 | O1P | 47.033 |

**Table S7.** Binding parameters of N-NTD and N-NTD-SR to different DNAs in 20 mM Bis-Tris pH 6.5.

| Protein | DNA | T (°C) | Kd (nM) | ΔH (kJ/mol) | ΔG (kJ/mol) | T·ΔS (kJ/mol) |
|---|---|---|---|---|---|---|
| N-NTD | | 15 | 9 ± 4 | | –44 ± 1 | 100 ± 2 |
| | **ssTRS(–)** | 25 | 4 ± 2 | 56 ± 1 | –47 ± 1 | 103 ± 2 |
| | | 35 | 2 ± 1 | | –51 ± 1 | 107 ± 2 |
| | | 15 | 45 ± 12 | | –41 ± 1 | 103 ± 1 |
| | **ssTRS(+)** | 25 | 19 ± 7 | 62 ± 1 | –44 ± 1 | 106 ± 1 |
| | | 35 | 8 ± 7 | | –48 ± 2 | 110 ± 2 |
| | | 15 | 6 ± 2 | | –45 ± 1 | –15 ± 2 |
| | **dsTRS** | 20 | 9 ± 3 | –60 ± 2 | –45 ± 1 | –15 ± 2 |
| | | 25 | 14 ± 4 | | –45 ± 1 | –15 ± 2 |
| | | 15 | 16 ± 8 | | –43 ± 1 | 68 ± 1 |
| | **ssNS(–)** | 25 | 11 ± 10 | 25 ± 1 | –45 ± 2 | 70 ± 2 |
| | | 35 | 8 ± 5 | | –48 ± 2 | 73 ± 2 |
| | | 15 | 17 ± 4 | | –43 ± 1 | 29 ± 2 |
| | **ssNS(+)** | 25 | 20 ± 4 | –14 ± 2 | –44 ± 1 | 30 ± 2 |
| | | 35 | 25 ± 6 | | –45 ± 1 | 31 ± 2 |
| | | 15 | 30 ± 8 | | –41 ± 1 | 1 ± 1 |
| | **dsNS** | 20 | 40 ± 11 | –42 ± 1 | –41 ± 1 | 1 ± 1 |
| | | 25 | 54 ± 12 | | –41 ± 1 | 1 ± 1 |
| N-NTD-SR | | 15 | 2 ± 1 | | –48 ± 1 | 109 ± 10 |
| | **ssTRS(–)** | 25 | 1 ± 2 | 61 ± 10 | –51 ± 5 | 112 ± 11 |
| | | 35 | 0.3 ± 1 | | –56 ± 8 | 117 ± 13 |
| | | 15 | 31 ± 6 | | –41 ± 1 | 87 ± 1 |
| | **ssTRS(+)** | 25 | 16 ± 7 | 46 ± 1 | –44 ± 1 | 90 ± 1 |
| | | 35 | 9 ± 7 | | –47 ± 2 | 93 ± 2 |
| | | 15 | 5 ± 2 | | –46 ± 1 | –99 ± 1 |
| | **dsTRS** | 20 | 14 ± 3 | –145± 1 | –44 ± 1 | –101 ± 1 |
| | | 25 | 38 ± 6 | | –42 ± 1 | –103 ± 1 |
| | | 15 | 36 ± 13 | | –41 ± 1 | 32 ± 1 |
| | **ssNS(–)** | 25 | 41 ± 6 | –9 ± 1 | –42 ± 1 | 33 ± 1 |
| | | 35 | 46 ± 6 | | –43 ± 1 | 34 ± 1 |
| | | 15 | 5 ± 4 | | –46 ± 2 | –61 ± 2 |
| | **ssNS(+)** | 25 | 24 ± 10 | –106 ± 2 | –43 ± 1 | –63 ± 2 |
| | | 35 | 94 ± 19 | | –41 ± 1 | –65 ± 2 |
| | | 15 | 89 ± 16 | | –39 ± 1 | 171 ± 1 |
| | **dsNS** | 20 | 35 ± 14 | 132 ± 1 | –42 ± 1 | 174 ± 1 |
| | | 25 | 14 ± 4 | | –45 ± 1 | 177 ± 1 |

**Table S8.** Affinity of interaction between N-NTD and DNAs estimated by CSP.

| Residue/Region | Kd (μM) | | | |
|---|---|---|---|---|
| | ssTRS(+) | ssTRS(–) | ssNS(+) | ssNS(–) |
| **N-terminal** | | | | |
| N47 | 8±12 | 4±5 | 19±11 | 5±3 |
| N48 | 4±3 | 3±1 | 16±6 | 6±2 |
| **β-sheet II (β1/β5)** | | | | |
| T54 | 1±2 | 0.1±0.8 | 11±10 | 11±7 |
| L56 | 4±3 | 5±5 | 22±9 | 1±4 |
| **Remote region** | | | | |
| D63 | 4±5 | 2±3 | 9±8 | 2±7 |
| K65 | 5±2 | 1.1±0.6 | 7±2 | 4±2 |
| F66 | 7±11 | 1±2 | 6±5 | 6±7 |
| Q163 | 1±2 | 3±2 | 18±6 | 12±7 |
| T166 | 4±1 | - | 11±2 | 6±2 |
| L167 | 5±4 | 0.3±0.6 | 21±7 | 1±1 |
| **Finger** | | | | |
| F90 | 13±7 | 7±5 | 18±13 | 1±2 |
| R95 | 1±4 | 0.1±0.5 | 1±2 | 4±3 |
| G96 | 3±9 | 1±5 | 24±18 | 5±6 |
| K102 | 1±3 | - | 21±11 | 3±2 |
| **Palm (β2/β3/β4)** | | | | |
| Y111 | 2±2 | 0.4±0.5 | 1±1 | 2.3±0.9 |
| **Thumb** | | | | |
| A152 | 0.7±0.7 | 0.4±0.4 | 8±3 | 2.5±0.6 |
| **C-terminal** | | | | |
| S176 | 3±2 | 1±5 | 6±7 | 1±3 |
| G178 | 2±2 | 0.1±0.5 | 5±4 | 9±12 |
| G179 | 2±4 | 2±5 | 3±5 | 2±5 |
| **All regions** | | | | |
| **Average** | 4±3 | 2±2 | 12±7 | 4±3 |
